# Impacts of ocean acidification and warming on post-larval growth and metabolism in two populations of the great scallop (*Pecten maximus* L.)

**DOI:** 10.1101/2022.12.02.518857

**Authors:** E. Harney, S.P.S. Rastrick, S. Artigaud, J. Pisapia, B. Bernay, P. Miner, V. Pichereau, Ø. Strand, P. Boudry, G. Charrier

## Abstract

Ocean acidification and warming are key stressors for many marine organisms. Some organisms display physiological acclimatisation or plasticity, but this may vary across species ranges, especially if populations are adapted to local climatic conditions. Understanding how acclimatisation potential varies among populations is therefore important in predicting species responses to climate change. We carried out a common garden experiment to investigate how different populations of the economically important great scallop (*Pecten maximus*) from France and Norway responded to variation in temperature and *p*CO_2_ concentration. After acclimation, post-larval scallops (spat) were reared for 31 days at one of two temperatures (13°C and 19°C) under either ambient or elevated *p*CO_2_ (pH 8.0 and pH 7.7). We combined measures of proteomic, metabolic, and phenotypic traits to produce an integrative picture of how physiological plasticity varies between the populations. The proteome of French spat showed significant sensitivity to environmental variation, with 12 metabolic, structural and stress-response proteins responding to temperature and/or *p*CO_2_. Principal component analysis revealed seven energy metabolism proteins in French spat that were consistent with countering ROS stress under elevated temperature. Oxygen uptake in French spat did not change under elevated temperature, but increased under elevated *p*CO_2_. In contrast, Norwegian spat reduced oxygen uptake under both elevated temperature and *p*CO_2_. Metabolic plasticity seemingly allowed French scallops to maintain greater energy availability for growth than Norwegian spat. However, increased physiological plasticity and growth in French spat may come at a cost, as French (but not Norwegian) spat showed reduced survival under elevated temperature.

**Summary Statement:** Juvenile scallops from France and Norway differ in their response to warming and acidification. French scallops show more physiological plasticity, adjusting their proteome and metabolism in order to maintain growth.

## 1. Introduction

Elevated atmospheric CO_2_ is a major driver of global climate change (Crowley & Berner 2001), causing surface temperatures to rise both on land and in the ocean (Hansen et al. 2006). Oceans act as a sink for more than a third of all anthropogenic carbon emissions (Sabine et al. 2004) leading to changes in marine carbonate chemistry and acidification of marine environments (Caldeira & Wickett 2003, Doney et al. 2009). Changes in temperature and *p*CO_2_ exert strong impacts on populations of ectothermic marine organisms (Pörtner 2002, Brierley & Kingsford 2009), especially those organisms that construct their shells from calcium carbonate (Ries et al. 2009). Furthermore, when experienced simultaneously, ocean acidification and warming (OAW) can result in synergistic or unforeseen effects (Pörtner & Farrell 2008, Todgham & Stillman 2013, Davis et al. 2013).

Phenotypic responses to OAW vary greatly amongst species (Kroeker et al. 2013, Scanes et al. 2014, Okazaki et al. 2017) and even within species (Morley et al. 2009, Pespeni et al. 2013a, Dam 2013, Calosi et al. 2017, Vargas et al. 2017). This variation may contribute towards phenotypic evolution, assuming that it has a heritable basis (Pespeni et al. 2013b, Dam et al. 2021). One of the key selective factors that could drive local adaptation is temperature variation along latitudinal gradients (Pereira et al. 2017). For example, towards lower latitude (warmer) range edges, populations may live close to their thermal limits (Pereira et al. 2017), and increases in temperature may induce poleward range shifts (Hale et al. 2017). Ectotherms from more thermally variable (temperate and boreal) latitudes may have greater thermal tolerance and ability to acclimatise than those from thermally stable (polar and equatorial) latitudes (Sunday et al. 2011). Furthermore, populations from higher latitude (colder) range edges may be less able to adjust their metabolism under elevated *p*CO_2_ (Calosi et al. 2017) and suffer reduced metabolism under combined stresses (Di Santo 2016). Yet, because metabolic rates increase exponentially with temperature, a given range of metabolic rates occupies a narrower range of temperatures in warmer climes (Payne & Smith 2017), which could allow for greater acclimatisation potential in cooler parts of the range. The fact that congeneric species can differ in how they adjust metabolism along their latitudinal distributions (Whiteley et al. 2011, Rastrick & Whiteley 2013) reinforces the importance of studying evolved differences in response to environmental variation.

Metabolism is therefore a key determinant of response to environmental variation among marine ectotherms. The ability to maintain oxygen supply is crucial for physiological performance (Byrne 2011, Pörtner 2012) and determining allocation of resources to competing energetic demands (Sokolova et al. 2012) with consequences for life history traits such as growth and fecundity. Chronic temperature stress frequently leads to elevated metabolic rates (Lefevre 2016), but can also result in acclimatisation and maintenance of ‘normal’ metabolic rates (Seebacher et al. 2010), or reduced metabolism (Anestis et al. 2008, Clark et al. 2013). Effects of increased *p*CO_2_ on metabolism also appear variable (Lefevre 2016) and may differ across life stages (Pörtner et al. 2010). In the long-term, calcifying organisms generally increase their metabolic rate in acidified conditions (Rastrick et al. 2018), or when warming and acidification are experienced simultaneously (e.g.: Matoo et al. 2013), but the physiological response may be dependent on the mechanisms and costs of maintaining acid-base status (Small et al. 2015).

Both **Error! Bookmark not defined.**temperatures and *p*CO_2_ charges are perceived and interpreted via a wide range of cellular signalling and metabolic pathways, which then facilitate acclimatisation via physiological plasticity (Seebacher et al. 2010, Pörtner 2012, Hurd et al. 2020). This acclimatisation is likely to play a key role in shaping tolerance to environmental stress (Seebacher et al. 2015), although it may also limit local adaptation (Sanford & Kelly 2011). Our understanding of the molecular mechanisms underlying acclimatisation has been aided by the development of high throughput ‘omics technologies (Mykles et al. 2010), including environmental proteomics (Tomanek 2011), which can reveal multifaceted responses to variation in the environment and climate stress. Relating proteomic responses to energetic trade-offs and in turn complex phenotypes (such as rates of growth, development, and survival) can provide clues about potential links between molecular responses and their fitness consequences (Artigaud et al. 2015, Harney et al. 2016, Timmins-Schiffman et al. 2020). Meanwhile, comparisons of congeners or different populations can reveal functionally adaptive patterns of protein abundance (Fields et al. 2012, Tomanek 2014). Integrating responses across molecular (e.g. proteomic), physiological, phenotypic, and population scales is necessary to best predict how species will respond to global climate change (Pörtner et al. 2006, Pörtner 2012).

It is generally accepted that molluscs are particularly at risk from ocean acidification and warming (Harvey et al. 2013, Kroeker et al. 2013), even though molluscan taxa differ in their susceptibility. Among bivalves, scallops may be more tolerant to acidification than oysters and mussels (Scanes et al. 2014), and appear to adjust their metabolism under combined stresses (Götze et al. 2020). Populations of great (or king) scallop *Pecten maximus* (L.) from temperate waters appear better able to maintain acid-base homeostasis than those from boreal waters (Schalkhausser et al. 2013, 2014), yet the molecular mechanisms that confer this tolerance and the phenotypic consequences of chronic exposure to elevated temperature and *p*CO_2_ remain poorly understood. Globally, scallop fisheries represent important economic resources, and successfully managing this resource requires a better understanding of how OAW will impact scallops at the population level (Rheuban et al. 2018). In the north-eastern Atlantic, *P. maximus* is an economically important species that is exploited along its native range, from Norway to Spain (Duncan et al. 2016). Interestingly, *P. maximus* from Norway appear to be genetically distinct from other European populations (Morvezen et al. 2015, Vendrami et al. 2019), and adults from the Bay of Brest in France and the North Sea near the western fjords of Norway display differences in growth phenotypes (Chauvaud et al. 2012) and proteomic profile (Artigaud et al. 2014b). Such differences may reflect environmental variation between the sites where these individuals were sampled. Mean sea surface temperature (SST) is higher in the Bay of Brest (13°C) than in the North Sea near the western fjords (8°C; Chauvaud et al. 2012). Furthermore *p*CO_2_ values are higher and more seasonally variable in the Bay of Brest (325-475 μatm) than in the North Sea near the western fjords (285-385 μatm; Salt et al. 2016, Omar et al. 2019). However, it is not clear whether the observed phenotypic and proteomic differences between these two populations are fixed or plastic, which is essential in determining whether these traits are heritable and could have an adaptive basis. Therefore, to improve our understanding of how *P. maximus* populations might respond to future climate change, we carried out common garden experiments in the lab using juvenile *P. maximus* (known as spat) from French and Norwegian populations. Understanding how changing environmental conditions affect sensitive early life stages is of particular importance, because these represent a bottleneck for population persistence (Byrne 2012). Spat were reared at three temperatures and two *p*CO_2_ concentrations (6 treatments) over a 5-week period and phenotypic responses were measured at multiple levels of biological organization, including protein abundance, oxygen consumption and growth in soft tissue and calcified structures.

## 2. Materials and Methods

### 2.1. *Production of* Pecten maximus *spat*

For the first few months after metamorphosis, post-larval *P. maximus* are commonly known as spat (although there is no further developmental transition before maturation, they are only referred to as juveniles after the first year; Christophersen et al. 2008). To test how spat respond to increased temperatures and *p*CO_2_, we carried out a common garden experiment at our experimental facilities. Both Norwegian and French spat were obtained from commercial scallop hatcheries (*Scalpro AS*: 5337, Rong, Vestland, Norway and *Écloserie du Tinduff*: Port du Tinduff 29470, Plougastel Daoulas, Brittany, France). At these hatcheries, adults collected in the wild are induced to spawn and offspring are reared from fertilized eggs through to early spat, before being transferred to sea cages to complete this phase of growth. Because of limitations on the availability of spat and differences in hatchery practices, Norwegian and French spat varied in their developmental history at the start of the experiment. Norwegian spat (offspring of approximately 60 adults sampled near Bergen, Norway) were approximately 7 months old and had yet to be placed in sea cages, while French Spat (offspring of approximately 30 adults sampled in the Rade de Brest, France) were 3 months old and had spent 2 weeks in sea cages.

Transport of approximately 3000 Norwegian spat from the hatchery in Rong, Norway to the experimental facility at Ifemer, Centre de Bretagne, France, followed the recommendations of Christophersen et al (2008) and took approximately 12 hours. Transport was carried out following submission and approval of an EU intra-trade certificate submitted via the TRACES platform. Spat were removed from their tanks and transferred to a cooled container (11°C) containing seawater-soaked absorbent paper for road transport to Bergen airport and air transport to Paris. On arrival in Paris, spat were transferred to a large (1000 L) tank containing seawater (maintained at 13°C in a refrigerated van). From here, they were transported to the experimental facility and transferred to tanks maintained at 13°C.

Approximately 9000 French spat were collected from sea cages near Sainte-Anne du Portzic, in the Bay of Brest, 6 days after the transport of Norwegian spat. We replicated the most stressful part of the transportation of Norwegian spat (emersion for approximately 6 hours) for French spat before introducing them to tanks.

### 2.2. Characterisation of experimental system and animal maintenance

Following transport or simulated transport, spat were transferred to six ‘raceway’ flow through tanks (100 L). For each population, spat were split among 18 mesh-bottomed trays (mesh 500 μm), held approximately 2.5 cm from the tank bottom by PVC supports, with populations kept separate initially (6 trays per raceway, 36 total trays). Each raceway drained into an independent header tank (30 L) containing an overflow. The rate of water renewal was regulated by gravity pressure between the input and overflow: filtered UV-sterilized sea water was supplied at a flow of approximately 90 mL min^-1^, leading to approximately one renewal per day. Header tanks also received a constant flow of microalgae (equal concentrations of *Tisochrysis lutea* and *Chaetoceros gracilis* at a final concentration of approximately 80 000 cells mL^-1^ in experimental tanks), which was supplied to two different header tanks from 10 L bottles via peristaltic pumps (two delivery tubes per pump, one for each header tank). Bottles of algae were replenished every 2 days.

From each header tank, a submerged pump supplied algae enriched seawater to a network of PVC pipes (with small holes drilled in them) overhanging each mesh-bottomed tray in the raceway at a rate of about 400 litres h^-1^or approximately 66 litres h^-1^ per tray. Hereafter we refer to each interconnected header tank and raceway as an experimental system, or system for short. The six systems were disinfected and rinsed on a weekly basis, with spat trays transferred to small tanks during cleaning (< 30 minutes per system). During cleaning, dead individuals and empty shells were removed from trays.

Spat were acclimated for 10 days (Norwegian spat) or 6 days (French spat) at 14.2 ± 0.5°C (ambient *p*CO_2_). Then, during an adjustment period of 6 days, temperatures were slowly changed in all six systems to reach treatment conditions (Fig. S1A), nominally 13, 16 and 19°C (two systems at each temperature). Temperatures in the 16 and 19°C systems were increased using resistance heaters placed in header tanks, temperatures in the 13°C systems were reduced by decreasing the temperature of in-flowing water. During acclimation and the initial part of the adjustment period, each raceway housed trays of a single population. On the penultimate day of the adjustment period, raceways were rearranged such that each contained an alternating sequence of Norwegian and French spat (three trays of each). The concentration of *p*CO_2_ was then elevated for one system at each temperature by bubbling CO_2_ through a CO_2_ reactor (JBL GmbH & Co. KG, Neuhofen, Germany) in the header tank. Elevated *p*CO_2_ was maintained by negative feedback based on a target pH of 7.7 (compared to 8.0 in untreated tanks), in line with end-of-the century predictions under rcp 8.5 (IPCC 2022). Experimental treatments were maintained for 31 days (from hereafter referred to as days 0-31; Fig. S1A-B). We hereafter use the nominal target temperatures and the abbreviations normCO_2_ (normal *p*CO_2_ / pH 8.0) and highCO_2_ (high *p*CO_2_ / pH 7.7) when referring to the different treatments (13-normCO_2_, 13-highCO_2_, 16-normCO_2_, 16-highCO_2_, 19-normCO_2_, and 19-highCO_2_).

Temperature was measured with a digital temperature probe and pH was measured using a WTW pH 340i fitted with a WTW SenTix 41 pH electrode (WTW GmbH, Weilheim, Germany). During acclimation and adjustment, conditions in header tanks were checked daily (excluding weekends), and during the experimental treatment, conditions were monitored twice daily (and once every two days during the weekend). Mean temperature and pH values for the six treatments are shown in table 1 and Fig. S1C-D.

**Table 1.** Mean (± standard deviation) environmental parameters during experimental treatments (days 0-31).

| Nominal<br>Treatment<br>(temp / pH) | Temperature *<br>(°C) | pH * | [HCO<sub>3</sub><sup>-</sup>] **<br>(μmol Kg<sup>-1</sup>) | DIC ***<br>(μmol Kg<sup>-1</sup>) | A<sub>T</sub> ***<br>(μmol Kg<sup>-1</sup>) | pCO<sub>2</sub> ***<br>(μatm) | Ω Calcite *** | Ω Aragonite<br>*** |
| --- | --- | --- | --- | --- | --- | --- | --- | --- |
| 13 / 8.0<br>13-normCO<sub>2</sub> | 13.37 ± 0.38 | 7.95 ± 0.06 | 2184 ± 24 | 2322 ± 33 | 2462 ± 53 | 680 ± 50 | 2.63 ± 0.29 | 1.68 ± 0.19 |
| 13 / 7.7<br>13-highCO<sub>2</sub> | 13.45 ± 0.37 | 7.70 ± 0.05 | 2206 ± 29 | 2317 ± 31 | 2356 ± 33 | 1307 ± 86 | 1.41 ± 0.11 | 0.90 ± 0.07 |
| 16 / 8.0<br>16-normCO<sub>2</sub> | 16.20 ± 0.60 | 7.93 ± 0.08 | 2180 ± 10 | 2324 ± 3 | 2473 ± 11 | 740 ± 56 | 2.79 ± 0.19 | 1.80 ± 0.12 |
| 16 / 7.7<br>16-highCO<sub>2</sub> | 16.20 ± 0.26 | 7.68 ± 0.05 | 2214 ± 16 | 2328 ± 17 | 2371 ± 22 | 1449 ± 61 | 1.48 ± 0.08 | 0.96 ± 0.05 |
| 19 / 8.0<br>19-normCO<sub>2</sub> | 19.12 ± 0.30 | 7.93 ± 0.06 | 2219 ± 23 | 2374 ± 21 | 2535 ± 16 | 808 ± 44 | 3.03 ± 0.12 | 1.97 ± 0.08 |
| 19 / 7.7<br>19-highCO<sub>2</sub> | 19.20 ± 0.39 | 7.65 ± 0.04 | 2241 ± 15 | 2360 ± 16 | 2401 ± 17 | 1660 ± 5 | 1.51 ± 0.02 | 0.98 ± 0.01 |

\* Temperature and pH were frequently measured (twice daily during weekdays, once per weekend).
\*\* Replicated bicarbonate ion ([HCO<sub>3</sub><sup>-</sup>]) measurements took place at two time points (days 23 and 31).
\*\*\* DIC (Dissolved Inorganic Carbon), A<sub>T</sub> (total alkalinity), pCO<sub>2</sub> (partial pressure of CO<sub>2</sub>), Ω Calcite and Ω Aragonite (calcite and aragonite saturation states)
were determined using the CO2SYS v2.1 macro (Pierrot et al. 2006).

At two points during the experiment (days 23 and 31) duplicated water samples from each experimental system were collected for salinity and alkalinity analyses. Salinity was determined using a refractometer and was found to be 36 PSU in all samples. Alkalinity was determined from bicarbonate ion [HCO_3_^-^] titration (analyses performed by Labocea laboratories, France). Bicarbonate concentration and pH were used to determine dissolved inorganic carbon (DIC) concentration. DIC, temperature, pH and salinity values were entered into the CO_2_SYS v2.1 macro (Pierrot et al. 2006) to calculate *p*CO_2_, total alkalinity, and saturation states of calcite and aragonite. The calculation was based on constants from Cai and Wang (Cai & Wang 1998) fitted to the NBS pH scale. Mean (± SD) carbonate chemistry conditions are shown in table 1.

### 2.3. Analysis of survival

Survival of spat in each tray was estimated by counting the number of individuals present in photos taken during the experimental treatment. By taking photos, we reduced the handling time and stress for the spat. On days 3, 9, 16, 24 and 31 (approximately once per week) photographs were taken of each of the 36 trays. To have sufficient image resolution, three photographs were taken of each tray and were stitched together afterwards using *GIMP* (GIMP development team 1997). All shells in stitched composites (apart from clearly empty shells) were counted using ImageJ software (Rasband 1997). Survival analysis was carried out with the *coxme* package (Therneau 2020) in the *R* statistical language (R Development Core Team 2019), with tray as a random factor to account for variation between trays within each experimental system. The significance of population and environmental variable effects on survival were tested with Wald chi-square tests using the *Anova* function from the *car* package (Fox & Weisberg 2011). Because *coxme* does not allow more than two dependent variables to be tested simultaneously, we first tested for differences between French and Norwegian scallops, before comparing the effects of temperature and *p*CO_2_ (plus their interaction) on survival for each population separately. To reduce the effect of sample size differences on statistical power, a random number of French spat, equivalent to the smaller number of Norwegian spat, were included in the survival analysis. Spat (2-6 individuals per tray) were sampled near-daily during the adjustment period and weekly during the experimental treatment for a separate experiment (unpublished data). These individuals were ‘left censored’ for the purposes of the survival analysis.

### 2.4. Analysis of whole organism phenotypes

At the end of the experiment (day 31), 20-40 individuals were removed from each tray and preserved in 95% ethanol at 4 °C. Three primary traits of shell size, shell weight and soft tissue weight were measured for between 8 and 15 individuals from each tray (mean = 12). Shell height is one of the most-commonly measured morphological phenotypes in bivalves. Bivalve shells grow by marginal accretion and changes in shell height provide an accurate measure of individual growth. Consistent patterns of accretion in the flat valves of scallops allows the estimation of growth over fine temporal scales (Chauvaud et al. 2012). Soft tissue dry weight (dry body weight) and total shell dry weight (total shell weight) are measures of investment in these two compartments. We also used these measures to estimate the condition index (CI: soft tissue dry weight / shell dry weight ratio), which encapsulates the difference in resource allocation to these compartments (Lucas & Beninger 1985). Spat were dissected and soft tissue was dried at 75 °C for 24 hours while both the left valve (flat) and right valve (curved) were air dried for at least 24 hours. The dry bodies, flat and curved valves were then weighed to the nearest 0.0001 g using a digital balance (Mettler Toledo).

Multiple high-resolution images of the flat valve at a range of focal depths were obtained using an AxioCam MRC 5 linked to a SteREO Lumar.V12 stereomicroscope (Carl Zeiss) equipped with a motorized stage: the resultant photomosaics were then assembled using AxioVision 4.9.1 software (Carl Zeiss). From these images shell height was measured using ImageJ software. These images were also used to estimate height at the beginning of the experiment (following transport), as growth in the controlled conditions of the experimental facility could be clearly associated with an alteration in the colour of newly calcified shell. Due to the considerable variation in size among scallops from both populations, initial shell height was included as a covariate in whole organism phenotypic analyses. Although Lucas and Beninger (1985) recommend the use of cubed height as a covariate for mass measures (such that variation scales in the same number of dimensions), we found initial height alone (not raised to a power) better accounted for covariation in the data. Phenotypic measures and ratios were plotted against these initial measures as part of an inspection for outliers. Four out of 431 samples were removed due to at least one trait showing extreme outlier values. Quantile-quantile plots were assessed to determine probability distributions. Dry body weight and total shell weight were subsequently log transformed to ensure normality. For all analyses, populations were analysed separately because of the strong differences in initial sizes.

The dependency of whole organism phenotypes on temperature, *p*CO_2_, and initial shell dimensions, as well as their 2-way and 3-way interactions, was assessed using linear mixed-effects models in the *lme4* package in *R* (Bates et al. 2015); tray was included as a random effect. Backwards stepwise term deletion was used to test the importance of interactions and main effects. Statistics were obtained from minimal models fitted with restricted maximum likelihood and *P*-values were obtained using the *lsmeans* package (Lenth 2016). When either temperature or the interaction of temperature and *p*CO_2_ were significant, contrasts between groups were evaluated with pairwise post-hoc tests using the *emmeans* package (Lenth 2020). For the interaction between temperature and *p*CO_2_, temperature differences at a given *p*CO_2_ and *p*CO_2_ differences at a given temperature were considered. Phenotypic responses to temperature and *p*CO_2_ were plotted using effect size plots in the *jtools* package (Long 2020) based on fully parameterised linear models (temperature, *p*CO_2_, initial shell dimensions, and all interactions). These account for covariate variation (including interactions), include confidence intervals, and can be mean centred. Although they do not account for random effect variance, these plots provide an intuitive means of visualising these data when combined with mixed-effects model statistics.

### 2.5. Analysis of metabolic rates

For both populations, the effect of temperature and *pCO_2_* on oxygen consumption ṀO_2_, used as a proxy for metabolic rate, was assessed (Rastrick et al. 2018) using randomly selected individuals from highest and lowest temperature treatments (13-normCO_2_, 13-highCO_2_, 19-normCO_2_, 19-highCO_2_). On day 27 of the experiment, three spat from each tray (nine per population/treatment combination, n = 72) were selected for metabolic analyses.

Spat were placed in individual stop-flow respirometers (volume 100ml) supplied with the same sea-water as the respective treatments. Animals were allowed 1 h to recover from handling and regain natural ventilatory behaviour before the flow to each chamber was stopped and the decreases in % oxygen saturation continuously measured using an optical oxygen system (Oxy-10mini, PreSense; labquest 2, Vernier; Rastrick and Whiteley, 2011; Rastrick 2018). The incubation period was 5 h, during which time, oxygen levels of the seawater did not fall below 70% (% air saturation) to avoid hypoxic conditions. A blank chamber with no animal was used to control for the background respiration in the seawater. The decrease in oxygen (% air saturation) within each chamber was converted to oxygen partial pressure (*p*O_2_) adjusted for atmospheric pressure and vapour pressure adjusted for relative humidity (measured using a multimeter; Labquest 2, Vernier). This decrease in *p*O_2_ was converted to concentration by multiplying by the volume of the chamber, minus the animal volume, and the oxygen solubility coefficient adjusted for the effect of temperature and salinity (Benson and Krause, 1984). Values were standardised to individual dry weight and expressed as μmol O_2_ h^-1^ mg^-1^ ± SEM. At the end of these metabolic experiments, the 72 individuals were sacrificed and dissected. Soft tissue was dried and weighed as described for the phenotypic analyses.

ṀO_2_ values were tested for normality. Although residuals approximated a normal distribution among French spat, they deviated from normality for Norwegian spat. Consequently, the *raov* function from the package *Rfit* (Kloke & McKean 2012) was used to provide rank-based estimations of linear models. We initially included population, temperature, *p*CO_2_ and all possible interactions in this model. However, to facilitate interpretation of the effects of temperature, *p*CO_2_ and their interaction, we also fitted models for each population separately. We used Benjamini-Hochberg-adjusted pairwise Wilcoxon tests to identify differences when the interaction was significant.

### 2.6. Analysis of the proteome

Spat for proteomic analyses were collected on day 31 from each tray in the 13-normCO_2_, 13-highCO_2_, 19-normCO_2_, and 19-highCO_2_ treatments. For each tray, two samples (each containing a pooled sample of two whole individuals) were flash frozen in liquid nitrogen (48 samples total) and stored at −80°C until analysis. Samples were homogenised by bead beating at 4°C in 500 μl Tris-HCl lysis buffer (100 mM, pH 6.8) containing 1% Protease Inhibitor Mix (GE Healthcare). A full and detailed description of the protocol for 2-dimensional gel electrophoresis (2-DE) and mass-spectrometry of protein samples can be found in Harney et al. (2016), but is described here briefly. Homogenised samples were centrifuged and solubilised proteins from the interphase were quantified using a D_C_ (detergent compatible) protein assay in a micro-plate reader. Then, 800 μg of protein were precipitated and desalted using a 1:1 ratio of sample to TCA/acetone (20% TCA). The supernatant was discarded, and pellets were neutralised by adding Tris-HCl/acetone (80% acetone) containing bromophenol blue as a pH indicator. Pellets were centrifuged once again and air-dried, before being rehydrated in Destreak rehydration solution (GE healthcare) containing 1% IPG (immobilised pH gradient) buffer (pH 4-7). After one hour, samples were ready for isoelectric focusing (IEF) on the IPGphor3 system (GE healthcare). After IEF, IPG strips were bathed in a rehydration solution (50 mM Tris–HCl pH 8.8, 6 M urea, 30% glycerol, 2% SDS and 0.002% Bromophenol Blue) for two 15 min periods, first with 10 mg/ml dithiothreitol, and then in the same solution containing 48 mg/ml iodoacetamide. Strips were then deposited on a 15cm × 15cm lab-cast SDS-PAGE gel containing 12% acrylamide and migrated. Protein spots were stained with Coomassie Blue (PhastGel, GE Healthcare). Gels were bleached with baths of H2O/methanol/acetic acid (70/30/7) and photographed using G:BOX (SynGene). The 32 clearest gels were taken forward for analysis (4 per population per treatment) and protein spots in the resulting images were aligned using Progenesis SameSpots v3.3 software (Nonlinear Dynamics, Newcastle upon Tyne, UK) and then manually verified. The effects of population, temperature, and pH were evaluated by running ANOVAs for each spot (combined population analysis). Due to the large number of tests involved, *P* values were adjusted using the false discovery rate (FDR), and fold change values were determined. Proteins which differed significantly in abundance between the populations, or between temperature or *p*CO_2_ treatments (FDR ≤ 0.05) were excised from the gels and analysed using mass spectrometry.

Gel pieces were first washed in 50 mM ammonium bicarbonate (BICAM), and then dehydrated in 100% acetonitrile (ACN). Gel pieces were vacuum-dried, rehydrated with BICAM containing 0.5 μg MS-grade porcine trypsin (Pierce Thermo Scientific), and incubated overnight at 37°C. Peptides were extracted from the gels by alternatively washing with 50 mM BICAM and ACN, and with 5% formic acid and ACN. Between each washing step, the supernatants from a given gel piece were pooled and finally concentrated by evaporation using a centrifugal evaporator (Concentrator 5301, Eppendorf).

Mass spectrometry (MS) experiments were carried out on an AB Sciex 5800 proteomics analyzer equipped with TOF/TOF ion optics and an OptiBeamTM on-axis laser irradiation with 1000 Hz repetition rate. The system was calibrated immediately before analysis with a mixture of Angiotensin I, Angiotensin II, Neurotensin, ACTH clip (1–17), and ACTH clip (18–39), showing that mass precision was above 50 ppm. After tryptic digestion, dry samples were resuspended in 10 μL of 0.1% TFA. A 1 μL volume of this peptide solution was mixed with 10 μL of 5 mg/mL α-cyano-4-hydroxycinnamic acid matrix prepared in a diluent solution of 50% ACN with 0.1% TFA. The mixture was spotted on a stainless steel Opti-TOF 384 target; the droplet was allowed to evaporate before introducing the target into the mass spectrometer. All acquisitions were taken in automatic mode. A laser intensity of 3400 was typically employed for ionizing. MS spectra were acquired in the positive reflector mode by summarizing 1000 single spectra (5 × 200) in the mass range from 700 to 4000 Da. Tandem mass spectrometry (MS/MS) spectra were acquired in the positive MS/MS reflector mode by summarizing a maximum of 2500 single spectra (10 × 250) with a laser intensity of 4200. For the MS/MS experiments, the acceleration voltage applied was 1 kV and air was used as the collision gas. Gas pressure was set to medium. The fragmentation pattern was used to determine the sequence of the peptide.

Database searching was performed using the MASCOT 2.4.0 program (Matrix Science). A custom database consisting of an EST database from a previous study was used (Artigaud et al. 2014c) and a compilation of the Uniprot database with *Pecten maximus* as the selected species. The variable modifications allowed were as follows: carbamidomethylation of cystein, K-acetylation, methionine oxidation, and dioxidation. “Trypsin” was selected as enzyme, and three miscleavages were also allowed. Mass accuracy was set to 300 ppm and 0.6 Da for MS and MS/MS mode, respectively. Protein identification was considered as unambiguous when a minimum of two peptides matched with a minimum score of 20. False discovery rates were also estimated using a reverse database as decoy.

As well as carrying out analysis of variance for the two populations combined, we also ran separate analyses of variance for each population. The overall effect of temperature, *p*CO_2_ and their interaction on protein abundance in each population were tested through permutational multivariate analysis of variance (Permanova) using the *adonis2* function in *vegan* (Oksanen et al. 2019). Then separate ANOVAs were fitted for each protein considering the effects of temperature, *p*CO_2_ and their interaction, with *P* values adjusted using FDR. For all proteins with significant environmental effects (FDR < 0.05), differences between the four treatments were quantified with post-hoc tests in *emmeans* (Lenth 2020).

To provide a clearer view of population responses to environmental variation, we ran additional exploratory and statistical analyses for each population separately using differentially abundant and successfully annotated proteins from the combined population proteomic analysis. We initially looked for correlations among proteins using principal component analysis (PCA), carried out in *R* using the packages *FactoMineR* (Lê et al. 2008) and *factoextra* (Kassambara & Mundt 2020), with spot size data scaled to unit variance. Correlations between proteins were identified by high loadings values (> 0.65 or < −0.65) of these variables onto principal components, which were visualised with vector plots. Differences between treatments were then visualised with individual coordinate plots and 95% confidence ellipses (the multidimensional space in which we expect to find the mean 95% of the time, given the underlying distribution of the data).

## 3. Results

### 3.1 Differences in survival

Survival was significantly higher among Norwegian spat than French spat (*χ*^2^ = 154.22, df = 1, *P* < 0.0001). For Norwegian spat, temperature, *p*CO_2_, and their interaction did not affect survival. For French spat, *p*CO_2_ did not affect survival (as a main effect or through its interaction with temperature); however, temperature did have a significant effect (*χ*^2^ = 18.79, df = 1, *P* < 0.0001), with mortality increasing with temperature. Survival curves are shown in Figure 1.

**Fig. 1.**
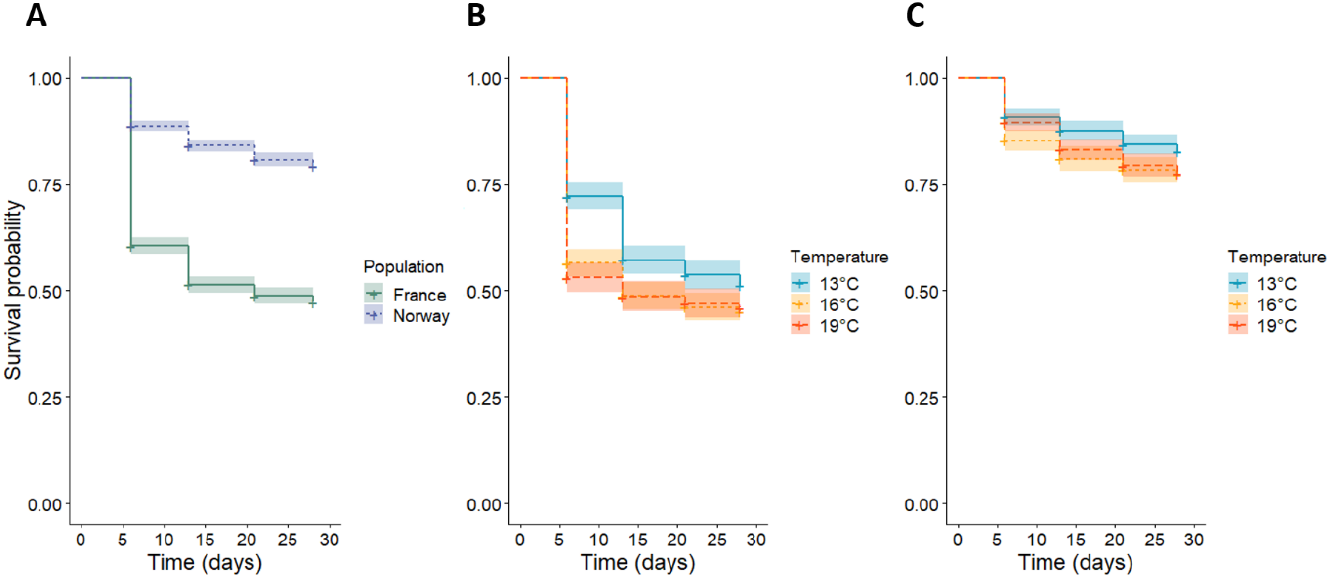
Results of survival analyses. **A**) French (green, n = 2556) and Norwegian (blue, n = 2556) spat survival. French spat had significantly lower survival compared to Norwegian spat (*χ*^2^ = 154.22, df = 1, *P*< 0.0001). **B**) Among French spat, increasing temperature (13, *n* = 829; 16, *n* =922; 19, *n* = 805) resulted in reduced survival (*χ*^2^ = 18.79, *P* < 0.0001). **C)** Among Norwegian spat (13, *n* = 882; 16, *n* =759; 19, *n* = 915), temperatures effects on survival were not significant (*χ*^2^ = 2.29, *P* = 0.319)

### 3.2. Variaition in whole-organism phenotypes

For the three primary traits of shell height, dry body weight and total shell weight, French and Norwegian spat differed markedly in their responses to *p*CO_2_ and temperature, although the effect of initial height was always highly significant (*P* < 0.0001). Among French spat, none of the primary traits responded strongly to temperature (table 2; Fig. 2A, 2C and 2E), and while elevated *p*CO_2_ had a positive effect on dry body weight (*F* = 4.92, df = 1, *P* = 0.041), it did not influence shell height or total shell weight. On the other hand, *p*CO_2_ effects were much stronger among Norwegian spat, where they interacted with initial height and temperature (table 2). For shell height, the *p*CO_2_ x temperature interaction was significant during the model selection process (in which ML estimates were used; *F* = 5.40, df = 2, *P* = 0.014), but the interaction was not significant once optimal models were refitted using REML estimates (*F* = 3.71, df = 2, *P* = 0.055). Yet the fact that the effects of temperature and *p*CO_2_ appear similar for all three primary traits in Norwegian spat (Fig. 2B, 2D and 2F) suggests that the *p*CO_2_ x temperature interaction for shell height, though weak, may be biologically meaningful. Thus, we report the *p*CO_2_-dependent temperature contrasts and temperature-dependent *p*CO_2_ contrasts for all three traits in table S1. At 19°C, elevated *p*CO_2_ had a significant negative effect on both shell height (*t* = 2.68, df = 11.9, *P* = 0.020) and dry body weight (*t* =2.42, df = 11.6, *P* = 0.033), and the effect was marginally non-significant for total shell weight (*t* = 2.12, df = 11.7, *P* = 0.056). In contrast, at 13°C, elevated *p*CO_2_ resulted in a greater total shell weight (*t* = −2.24, df =12.8, *P* = 0.043). Furthermore, at elevated *p*CO_2_ there was a decrease in dry body weight at 19°C relative to 13°C (*t* = 3.75, df 12.0, *P* = 0.007), and at normal *p*CO_2_ there was an increase in total shell weight at 16°C relative to 13°C (*t* = - 2.664, df 13.2, *P* = 0.0474).

**Fig. 2.**
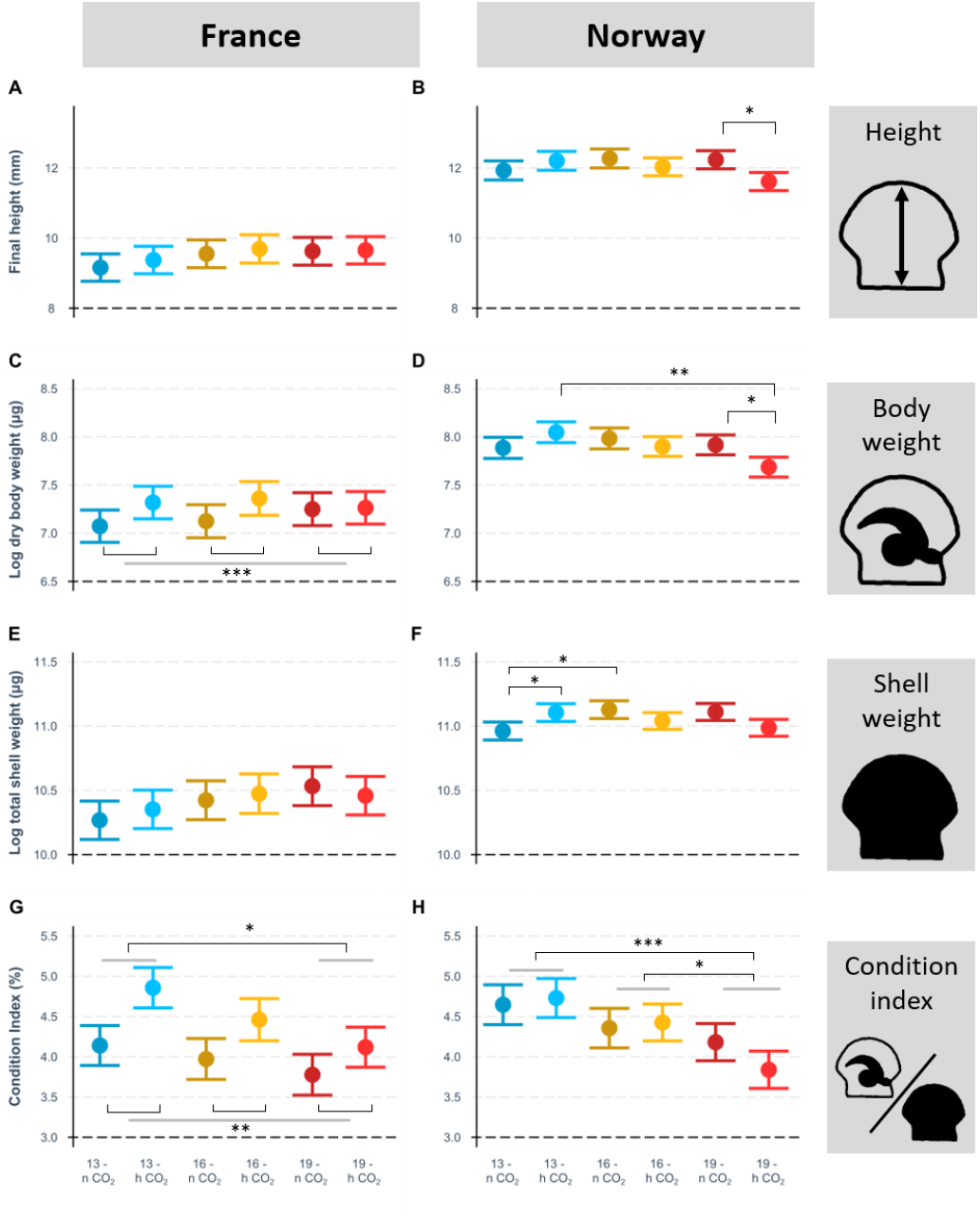
Effect of temperature and pH on French and Norwegian scallop whole organism phenotypic traits: final shell height (**A**, **B**), dry body weight (**C**, **D**), total shell weight (**E**, **F**) and condition index (**G**, **H**). Values are mean-centred model estimates (± c.i.) derived from linear models considering initial height, temperature, pH, and all their interactions. Effect/interaction significance was determined by term deletion and model comparison, and estimated marginal means were used to determine significant temperature and temperature x *p*CO_2_ contrasts. Simple brackets show temperature x *p*CO_2_ interactions, brackets linking grey bars show temperature effects (**G**, **H**), and grey bars linking brackets show *p*CO_2_ effects (**C**, **G**). For French spat, *n* = 36, 34, 35, 35, 33, 37; for Norwegian spat, *n* = 35, 35, 34, 39, 38, 40. All statistical tests and *P* values are shown in table 2.

**Table 2.**
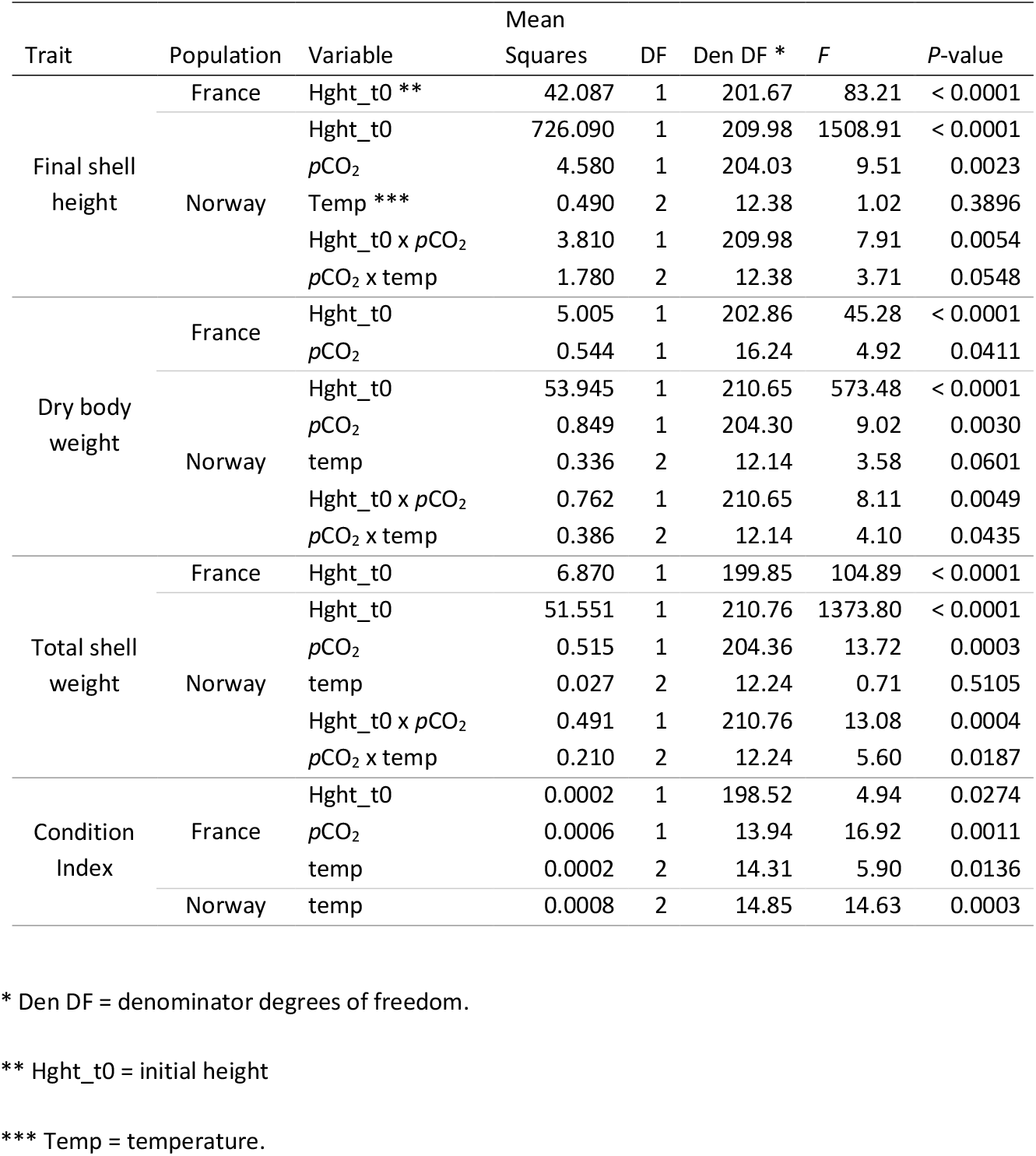
Summary table of effects and interactions of temperature, *p*CO_2_ and initial height on organismal phenotypes in minimal models following backwards stepwise term-deletion.

Although condition index (CI) was based on the ratio of two of the primary traits (dry body weight over total shell weight), it revealed new effects that were not identified from analyses of primary traits. Specifically, analysis of CI identified a significant temperature effect in both French (Fig. 2G; *F* = 5.90, df = 2, *P* = 0.014) and Norwegian spat (Fig. 2H; *F* = 16.63, df = 2, *P* < 0.001), with CI declining as temperature increased, particularly when comparing 13°C and 19°C treatments (table 2, table S1). CI also responded positively to elevated *p*CO_2_ in French spat (*F* = 16.92, df = 1, *P* = 0.001), mirroring the result found in dry body weight. Furthermore, initial height was not a significant covariate in explaining CI for Norwegian spat and had a weaker effect than temperature and *p*CO_2_ among French spat (*F* = 4.94, df = 1, *P* = 0.027).

### 3.3. Metabolic rate differences

French spat had higher average oxygen consumption (ṀO_2_) than Norwegian spat (91.98 μmol O2 h^-1^ mg^-1^ compared to 78.20 μmol O2 h^-1^ mg^-1^); however, a three way interaction between population, *p*CO_2_ and temperature (*F* = 7.79, df = 1, *P* < 0.0076) suggested the environmental effects differed strongly between populations, and models were refitted for each population separately. Among French spat, oxygen consumption (ṀO_2_) was significantly higher in elevated *p*CO_2_ treatments (*F* = 41.67, df = 1, *P* < 0.0001), but there was no effect of temperature (*F* = 0.44, df = 1, *P* = 0.516), nor any interaction between *p*CO_2_ and temperature (*F* = 0.51, df =1, *P* = 0.483). Among Norwegian spat, the interaction between *p*CO_2_ and temperature was significant (*F* = 5.77, df = 1, *P* = 0.025): pairwise post-hoc Wilcoxon rank sum tests confirmed that ṀO_2_ was significantly elevated (*P* < 0.05) in the 13-normCO_2_ treatment compared to the other treatments. ṀO_2_ differences between treatments are summarised in Fig. 3.

**Fig. 3.**
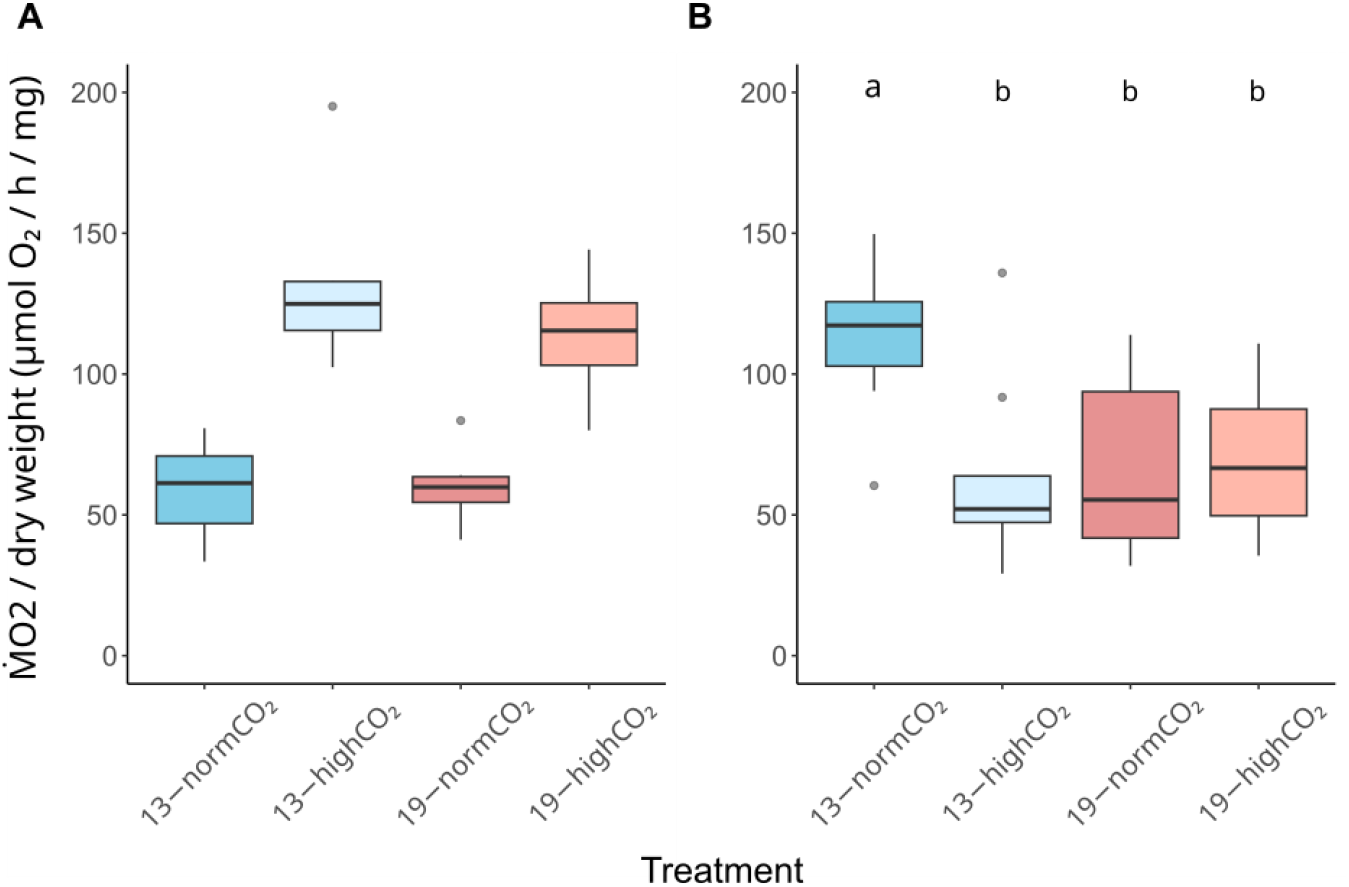
Weight-corrected oxygen consumption (ṀO_2_) in French and Norwegian spat after 5 weeks at experimental temperature and *p*CO_2_. French spat (A) displayed increased oxygen consumption under elevated *p*CO_2_ (*F* = 41.67, *P* < 0.0001; *n* = 7, 6, 6, 8). Among Norwegian spat (B), there was a significant interaction between elevated temperature and *p*CO_2_ (*F* = 5.77, *P* = 0.025; *n* = 7, 8, 6, 6). Post-hoc tests (letters a, b) revealed that increases in temperature and/or *p*CO_2_ resulted in reduced oxygen consumption.

### 3.4 Differential accumulation of proteins: combined population analysis

We identified 279 proteins common to all gels using SameSpots. Of these, 103 differed significantly (FDR < 0.05) between populations (n = 87), temperature treatments (n = 17) or *p*CO_2_ treatments (n = 3); three differed according to two or more of these variables. Following mass spectrometry of these proteins, 79 were identified based on comparison with the protein database: 71 differed according to population, 8 differed according to temperature and 2 differed according to *p*CO_2_ (two proteins differed according to two variables). These proteins are presented in table S2 and figure 4. Of the 79 proteins, 23 were highly differentially accumulated (|FC| > 2) and 33 were moderately differentially accumulated (|FC| > 1.5). Two proteins annotated as ‘uncharacterised’ were further investigated using nucleic acid homology searches. Spot 447 (*Mizuhopecten yessoensis* locus 110464099 showed strong amino acid similarity (65.14%, E = 7e-101) to the *Crassostrea gigas* cytoskeletal protease kyphoscoliosis peptidase (KY), while spot 468 (*Mizuhopecten yessoensis* locus 110453073) contained a conserved domain with significant similarity (interval 47-194, E = 4.19e-07) to von Willebrand factor A domain (vWA), an extracellular glycoprotein.

**Fig 4.**
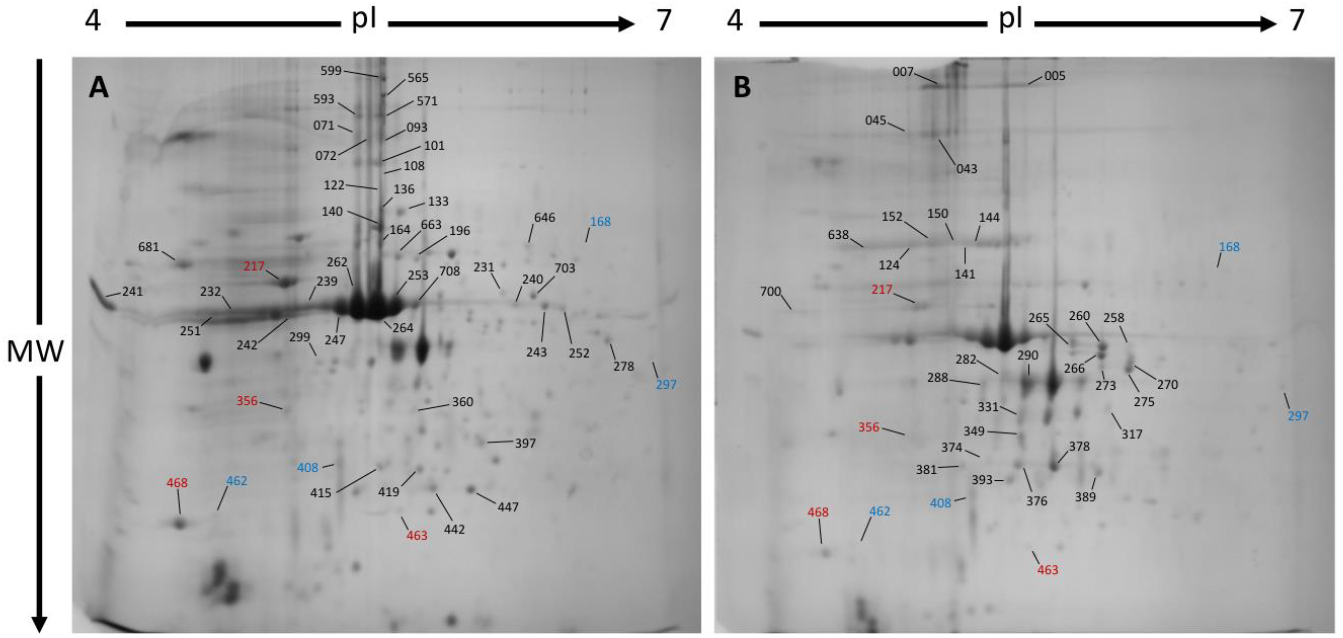
Representative annotated 2-DE gel images of French (A) and Norwegian (B) spat proteomes at 19°C and ambient *p*CO_2_. Proteins that were significantly more abundant in French scallops appear in (A); proteins that were significantly more abundant in Norwegian scallops appear in (B). Proteins with temperature-dependent abundance appear in both (A) and (B). Protein spots that were more abundant at 19°C are coloured red, those that were more abundant at 13°C are coloured blue. MW = molecular weight, pI = isoelectric point.

Seventy-one of the 79 proteins differed significantly between populations, and the majority of these (43) were different isoforms of actin: 24 isoforms were elevated among French spat, and 19 were elevated among Norwegian spat. Actin isoforms were spread widely across the 2-DE gel, and those that were more abundant in Norwegian spat generally had a higher pH and/or higher molecular weight. Another group of structural proteins which differed between the populations were motor proteins: 8 isoforms of myosin and 2 of paramyosin showed higher accumulation in Norwegian spat compared to French spat.

Of the 26 remaining proteins, the majority (18/26) differed between populations, and almost all of these (17/18) were more abundant in French spat than Norwegian spat: just one (gelsolin) showed higher accumulation in Norwegian spat. Nine proteins showed significant temperature and/or *p*CO_2_ effects in the combined population analysis, and five of these displayed fold changes greater than 1.5 (Fig. 5). The oxidative stress response protein manganese superoxide dismutase (MnSOD) was more abundant at 19°C (*F* = 14.90, *df* = 1, *FDR* = 0.008). Conversely, the extra cellular matrix protein ependymin-related 1 (EPDR), respiratory complex I (complex I) and the molecular chaperone peptidyl-prolyl cis-trans isomerase (PPIase) were more abundant at 13°C (EPDR: *F* =21.94, *df* = 1, *FDR* = 0.001; complex I: *F* =16.08, *df* = 1, *FDR* = 0.005; PPIase: *F* = 19.85, *df* = 1 *FDR* = 0.002). An isoform of paramyosin was more abundant at elevated *p*CO_2_ (*F* = 9.30, *df* = 1, *FDR* = 0.040) as well as being more abundant in Norwegian spat (*F* = 10.18, *df* = 1, *FDR* = 0.033).

**Fig 5.**
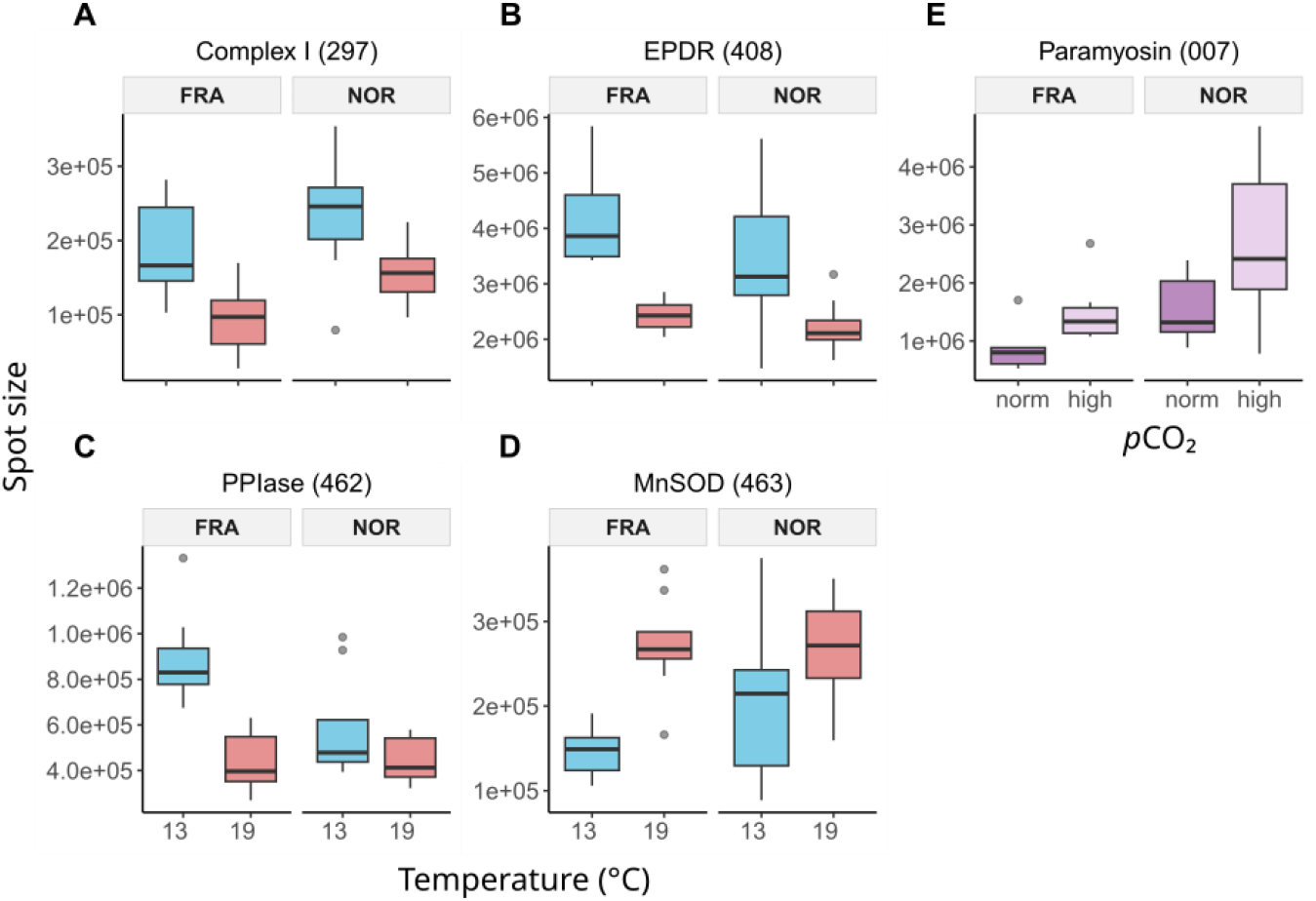
Proteins that differed significantly between temperature and *p*CO_2_ treatments in the combined population analysis. (FDR < 0.05, fold change > 1.5). For temperature responses (A-D) protein spots size from both *p*CO_2_ treatments are combined, and for *p*CO_2_ responses (E), protein spots from both temperatures are combined (in all cases *n* = 8). A) Complex I was more abundant at lower temperatures (*F* =16.08, *FDR* = 0.005), as were B) EPDR (*F* =21.94, *FDR* = 0.001) and C) PPIase (*F* = 19.85, *FDR* = 0.002). D) MnSOD was less abundant at lower temperatures (*F* = 14.90, *FDR* = 0.008), and E) a paramyosin isoform was more abundant at elevated *p*CO_2_ (*F* = 9.30, *FDR* = 0.040) and in Norwegian spat (*F* = 10.18, *df* = 1, *FDR* = 0.033).

### 3.5 Differential accumulation of proteins: separated populations analyses

The Permanova revealed that both temperature (*F* = 3.0444, df = 1, *P* = 0.002) and *p*CO_2_ (*F* = 3.3187, df = 1, *P* = 0.007) significantly influenced overall patterns of protein abundance in French spat, but not Norwegians spat (Temperature: *F* = 0.6211, df = 1, *P* = 0.721; *p*CO_2_: *F* = 0.4765, df = 1, *P* = 0.860). The interaction between temperature and *p*CO_2_ was not significant for either population. Among French spat, individual ANOVAs for the 79 proteins revealed 12 with potential temperature or *p*CO_2_ effects (FDR < 0.05; Fig. 6; table S3), six of which were also significant in the combined population analysis. Fold changes were greater than 1.5 in eight of these proteins. Although none of the 12 proteins showed a significant response to both temperature and *p*CO_2_ at our statistical threshold (FDR < 0.05), differing responses to elevated temperature and *p*CO_2_ were suggested by post-hoc tests (table S4). These showed that elevated temperature and *p*CO_2_ either had opposite effects that offset each other when combined (Fig. 6 A-G), or similar effects that exacerbated one another additively (Fig. 6 H-I). Conversely, among Norwegian spat only 11 proteins showed potential responses to temperature, *p*CO_2_ or their interaction (*P* < 0.05; Fig. S2), and none of these were significant following correction for multiple testing (FDR < 0.05).

**Fig 6.**
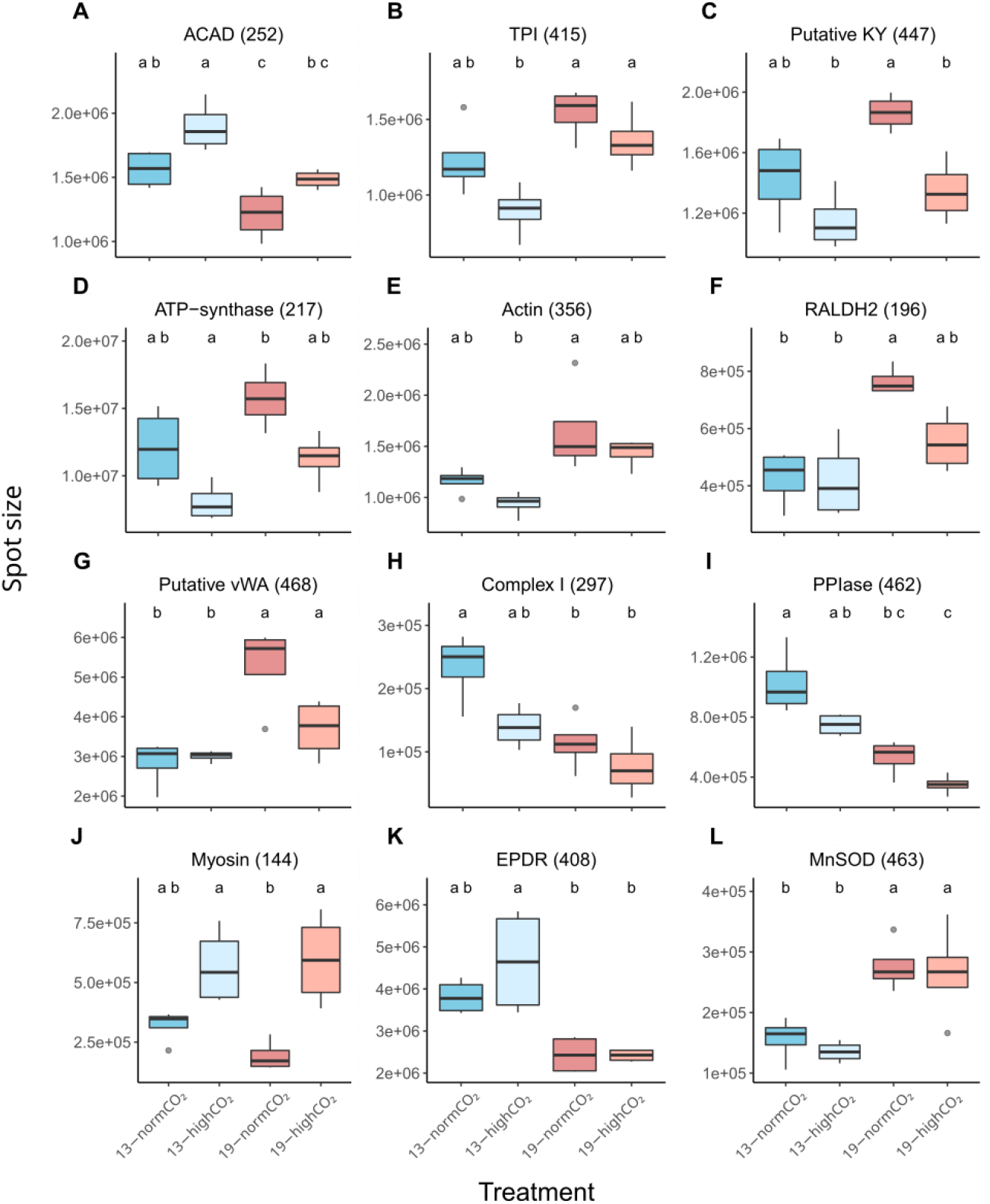
Proteins that differed significantly according to either temperature (A-B, E-L) or *p*CO_2_ (C-D) in French spat. (FDR < 0.05). Temperature effects (A-B, E-L) were more prevalent than *p*CO_2_ (C-D), effects, although post-hoc test results (indicated by a, b, c, etc. above plots) suggest that in many cases both were responsible for shaping protein abundance. ANOVA statistics are provided in table S3, post-hoc test statistics are provided in table S4. For all treatments in all proteins, *n* = 4.

The 12 environmentally dependent proteins in French scallops were ATP synthase, triosephosphate isomerase (TPI), medium-chain specific acyl-CoA dehydrogenase (ACAD), complex I, Retinal dehydrogenase 2 (RALDH2), MnSOD, PPIase, EPDR, an isoform of actin, an isoform of myosin, and putative isoforms of Kyphoscoliosis peptidase (KY) and von Willebrand factor type A (vWA). For these 12 proteins only one main effect, temperature (nine out of 12) or *p*CO_2_ (three out of 12), was significant at our stringent statistical cut-off (FDR < 0.05). Despite this, post-hoc tests indicated that both variables frequently had an impact on protein abundance (table S3). For seven proteins (Fig. 6A-G), the most significant difference in abundance occurred between 13-highCO_2_ and 19-normCO_2_ treatments, while the comparison of 13-normCO_2_ and 19-highCO_2_ did not differ significantly. This suggests that *p*CO_2_ and temperature had opposite effects that offset each other when both were elevated. For all these proteins (except ACAD, Fig. 6A), increasing temperature had a positive effect on abundance while elevated *p*CO_2_ had a negative effect. On the other hand, in two proteins (complex I and PPIase; Fig. 6H-I) the 13-normCO_2_ and 19-highCO_2_ treatments differed the most, suggesting that temperature and *p*CO_2_ both had additive negative effects on abundance.

### 3.6 Principal component analyses of protein abundance

To further explore correlations in protein abundance we carried out PCA using the 25 proteins that were not annotated as actin, myosin or paramyosin. Correlations were stronger among French spat, where the first two principal components (PC1 and PC2) together explained 54.7 % of the total variance (Fig. 7A) compared with 44.5% in Norwegian spat (Fig. 7B). Among French spat, 12 proteins had high loading values (> 0.65 or < −0.65) for PC1 (Fig. 7A), and treatments also showed separation along this axis in the individual coordinate plot (Fig. 7C). The confidence ellipse for 19-normCO_2_ treatment was associated with higher PC1 values than any other treatment, and the confidence ellipse for 13-highCO_2_ treatment was associated with lower PC1 values than ellipses for either 19-normCO_2_ or 19-highCO_2_. Transaldolase (TALDO), TPI_1, TPI_2, ATP synthase, RALDH1, RALDH2, glutathione S-transferase (GST), KY and vWA were positively correlated with PC1, while glucose-6-phosphate isomerase (GPI), ACAD, and an isoform of NADP-dependent isocitrate dehydrogenase (IDH_1) were negatively correlated with PC1. Among Norwegian spat, nine proteins had high loading values for PC1 (Fig. 7B), but there was no separation of treatments in the individual coordinate plot (Fig. 7D).

**Fig 7.**
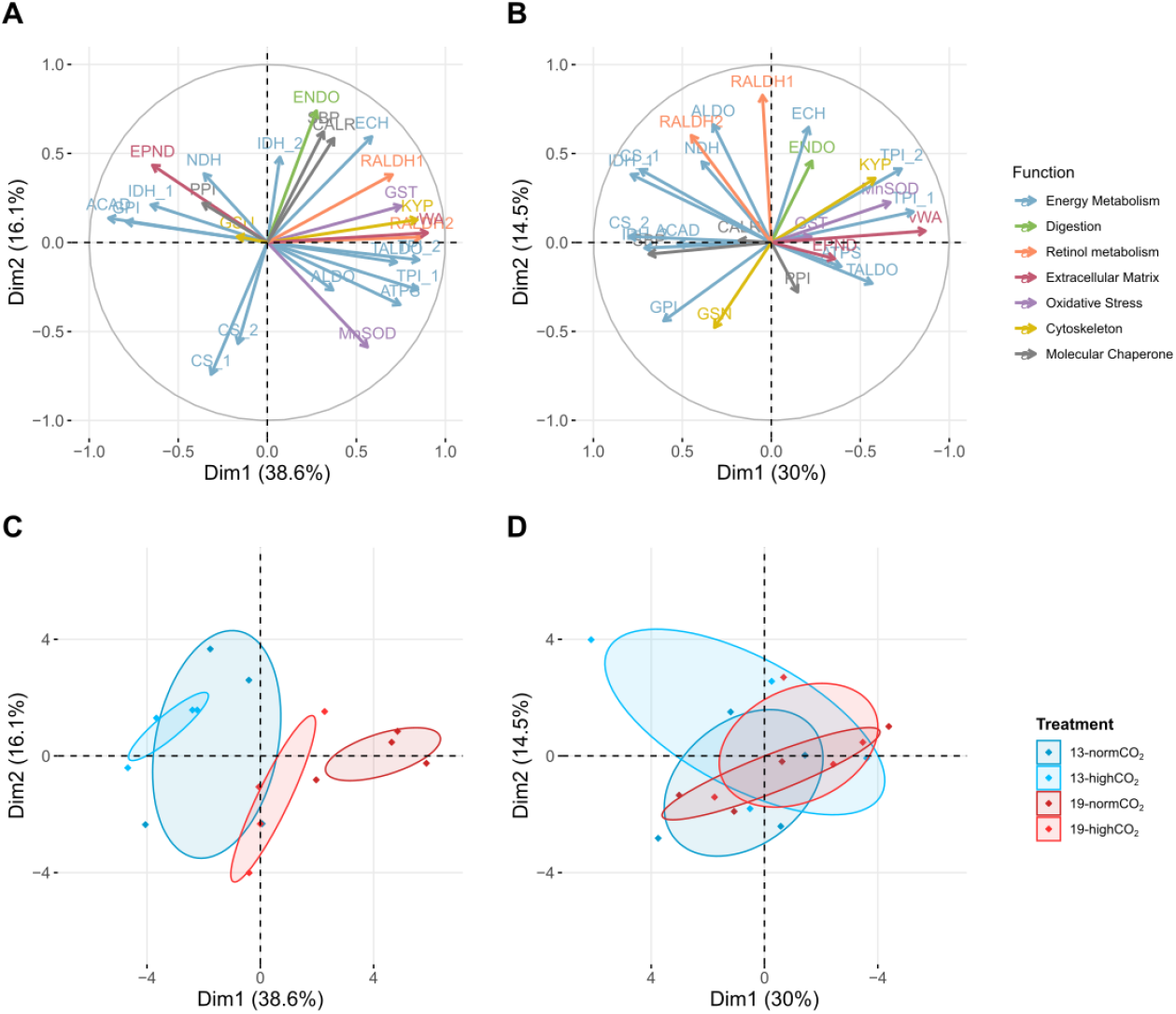
Vector and PCA plots of all proteins that were not actin, myosin or paramyosin. Vector plots show that patterns of protein correlation were stronger in French (A) than Norwegian (B) spat. Proteins are coloured according to their (putative) function. Most protein abbreviations are in text, and a full list is found in table S2. PCA plots show greater separation of treatments for French (C) than Norwegian (D) spat (95% confidence ellipses coloured according to temperature and *p*CO_2_ treatment). In both plots of Norwegian spat proteins (B and D), PC1 has been inverted to highlight similarities between the populations. For all treatments in all proteins, *n* = 4.

## 4. Discussion

### Phenotypic responses to environmental variation

Elevated temperature and *p*CO_2_ treatments both had significant antagonistic consequences for growth phenotypes in Norwegian scallops, with the strongest effects on growth experienced when both stresses were combined, a result that corresponds to other results in scallops (Artigaud et al. 2014a, Alma et al. 2020). In contrast, growth of French spat was less influenced by experimental treatments: the only primary phenotypic trait affected was dry body weight, which increased in elevated *p*CO_2_ treatments. However, elevated temperature resulted in greater mortality of French spat, suggesting a potential trade-off between growth and survival. While *P. maximus* adults have been shown to be fairly tolerant to warming and hypercapnic stresses (Götze et al. 2020), our results suggest spat may pay some costs under OAW: reduced survival in the French spat experiencing warming, and reduced growth in the Norwegian population experiencing acidification and warming.

Interestingly, condition index (CI; dry body weight divided by total shell weight), also revealed a clear positive effect of elevated *p*CO_2_ on CI in French spat. Similar results have previously been interpreted as seasonal shifts in patterns of resource allocation (Cameron et al. 2019), or subtle shifts in allocation to soft tissue and shell (Hiebenthal et al. 2013), but this result could also be an experimental artefact. It is possible that elevated *p*CO_2_ resulted in some microalgal growth (tanks were only semi-open) and increased food availability. However, if this were the case, only French scallops were able to exploit it, as neither dry body weight nor CI increased at elevated *p*CO_2_ in Norwegian spat. For scallops from both populations, CI declined with increasing temperatures in line with previously reported results from bivalves (Clark et al. 2013, Hiebenthal et al. 2013, Cameron et al. 2019, Pereira et al. 2020). This could be due to differences in the energetic costs of calcification compared with homeostasis: saturation states of aragonite and calcite increase at higher temperatures, potentially reducing the cost of calcification (Clark 2020, Clark et al. 2020). Furthermore, as ectotherms approach their upper thermal limits, aerobic scope is reduced (Pörtner & Farrell 2008, Sokolova et al. 2012), which can result in increased costs of basal metabolism and a relative decline in allocation of resources to soft tissue growth.

### Metabolic responses to increasing temperature and pCO_2_

As ectotherms approach upper thermal limits, they may reduce their metabolic rates (Anestis et al. 2008, Clark et al. 2013), which could help to explain why oxygen consumption of Norwegian scallops declined at higher temperatures. In contrast, oxygen consumption was not influenced by temperature in French scallops, suggesting that these spat successfully acclimated to experimental temperatures to maintain metabolism (Seebacher et al. 2010). Although oxygen consumption of French scallops did not vary according to temperature, there was a clear increase in oxygen consumption at elevated *p*CO_2_. This result mirrors findings in several other marine invertebrate species (Parker et al. 2012, Benítez et al. 2018, Harianto et al. 2021, Jiang et al. 2021), where increased ṀO_2_ in response to increased *p*CO2 appears to maintain cellular homeostasis. In contrast, Norwegian scallops reduced their oxygen consumption under elevated *p*CO_2_ irrespective of temperature. Given that this response occurred at 13°C, which Norwegian scallops experience during the summer, it could indicate alternative strategies for dealing with *p*CO_2_ variation between the populations, or that scallops from the Bay of Brest are better adapted to the more variable *p*CO_2_ levels that occur there (Salt et al. 2016). Genetic variation in ability to acclimate to elevated *p*CO_2_ has been observed previously among different selected lines of oysters (Parker et al. 2012) and mussels (Stapp et al. 2017), in which families tolerant of *p*CO_2_ variation were found to increase metabolic rates under elevated *p*CO_2_, while sensitive families did not. Given the strong and significant genetic differentiation between French and Norwegian scallop populations (Morvezen et al. 2015, Vendrami et al. 2019), it seems likely that higher temperatures and *p*CO_2_ in the Bay of Brest have led to an adaptive ability to acclimate to these conditions.

### Proteomic responses to environmental variation

Variation in the influence of temperature and *p*CO_2_ on metabolism was also detected in the proteome. While both populations responded to increased temperatures with increased abundance in the oxidative stress enzyme MnSOD and reduced abundance of the oxidative phosphorylation enzyme complex I (also known as quinone oxidoreductase and NADH dehydrogenase), the proteome of French spat showed far greater plasticity than that of Norwegian spat, again highlighting potential evolutionary differences between the populations. Among French spat, temperature effects were generally greater than *p*CO_2_ effects, but acidification frequently exerted a subtle effect in the opposite direction to heating, with increased temperature and *p*CO_2_ offsetting one another. This was emphasised by the PCA, in which increasing temperature had a positive effect on the first principal component, and increasing *p*CO_2_ had a negative effect. Four energy metabolism proteins (TALDO, TPI_1, TPI_2, and ATP-synthase) had positive loadings with PC1, and three energy metabolism proteins (GPI, IDH_1 and ACAD) had negative loadings. A decrease in GPI (the first enzyme involved in glycolysis) and concurrent increase in the pentose phosphate pathway (PPP) enzyme TALDO indicates a putative shift from the preparatory phase of glycolysis to the oxidative phase of PPP (Krüger et al. 2011). Furthermore, a greater abundance of TPI isoforms indicates that PPP metabolites are likely returning to the pay-off phase of glycolysis rather than being recycled between oxidative and non-oxidative branches of the PPP (as this would also require GPI; Krüger et al. 2011). TPI may be a particularly strong marker of this metabolic change because of its tendency to oligomerise at higher temperatures (Katebi & Jernigan 2014, Rodríguez-Bolaños et al. 2020). By directing carbon metabolism to the oxidative branch of the PPP, French scallops may be generating greater quantities of the reducing agent NADPH (Ralser et al. 2007, Stincone et al. 2015) to mitigate the increase in reactive oxygen species (ROS) production associated with metabolism at higher temperatures (Tomanek & Zuzow 2010). This idea is supported by a positive correlation in the abundance of the antioxidant GST, which is another critical component in managing ROS stress (Park et al. 2019).

Four other proteins (RALDH1, RALDH2, KY, and vWA) with putative roles in development, growth and biomineralisation also had high positive loadings for PC1 (increased abundance at high temperature, reduced abundance in elevated *p*CO_2_) in French scallops. Retinal dehydrogenases (RALDH1 and RALDH2) are involved in retinoic acid metabolism, which is associated with embryonic development, organ generation and homeostasis in vertebrates (Marlétaz et al. 2006), but are also known to affect development of the molluscan central nervous system (Carter et al. 2010, 2015). Kyphoscoliosis peptidase (KY) has previously been linked to molluscan stress responses (Chaney & Gracey 2011, Shiel et al. 2017, Blalock et al. 2020), but may also play a role in muscle growth (Shen et al. 2018). Finally the extracellular matrix protein vWA is likely to be involved in biomineralization (Funabara et al. 2014, Chandra & Vengatesen 2020, Clark et al. 2020). Beyond these results in French scallops, two other proteins with putative roles in biomineralization (EPDR and PPIase) were found to be more abundant at 13°C in the combined population analysis. EPDR has been directly implicated in molluscan biomineralization (Jackson et al. 2006, Marie et al. 2010, Miyamoto et al. 2013), while a subfamily of PPIases known as cyclophilins facilitate molluscan nacre formation (Jackson et al. 2010, Marie et al. 2013). Both temperature and *p*CO_2_ are known to have important effects on biomineralization processes and the crystalline structure of calcium carbonate (Fitzer et al. 2015), and our results provide some indications of the proteins that may underlie such changes.

### Population differences in cytoskeletal proteins

The structural proteins actin and myosin were among the most numerous in our study. Although actin has previously been implicated in bivalve physiological stress responses to both temperature (Tomanek et al. 2011) and *p*CO_2_ (Moreira et al. 2018), we found just one environmentally responsive isoform of actin which positively responded to increased temperature in French spat. Instead, actin isoforms differed substantially between the populations, with some more abundant in French scallops and others more abundant in Norwegian, echoing an earlier *in natura* comparison of adult *P. maximus* from these two populations (Artigaud et al. 2014b): results from our common garden approach provide more evidence that differences in proteomic abundance reflect divergent genetic backgrounds. Actin is abundant and multifunctional (Dominguez & Holmes 2011). Its density in our samples may be due to its presence in the scallop’s largest organ, the adductor muscle (Chantler 2006), where it aids with muscular contraction through its interaction with myosin. Indeed eight isoforms of myosin and two of the related protein paramyosin (both components of the adductor muscle; Chantler 2006) also showed strong population differentiation. However, unlike actin, these proteins were always more abundant in Norwegian spat. As they grow, scallop spat increasingly use their adductor muscle for swimming, which can lead to an increase in muscle condition (Kleinman et al. 1996). This could explain why Norwegian scallops (which were slightly older and larger at the start of the experiment) had elevated levels of these proteins. While no isoforms of myosin or paramyosin responded to temperature treatment, there was some evidence of *p*CO_2_ sensitivity in one isoform of myosin (more abundant at elevated *p*CO_2_ in French spat) and one isoform of paramyosin (more abundant at elevated *p*CO_2_ in the combined population analysis). Several recent studies from diverse marine invertebrates have linked increases in myosin and/or paramyosin transcript (Wäge et al. 2016, Bailey et al. 2017) and protein (Timmins-Schiffman et al. 2014, Zhao et al. 2020) abundance with the response to elevated *p*CO_2_. While the mechanism by which myosin and paramyosin abundance aids the response to acidification remains unclear, its presence in diverse taxa could suggest an evolutionary conserved physiological response.

### Integrating results across biological scales and conclusions

Drawing on phenotypic results from whole organismal, metabolic, and proteomic scales, we show clear differences in how French and Norwegian *Pecten maximus* spat respond to increases in temperature and *p*CO_2_. Although some proteomic and organismal responses were common to both populations, such as the increase in MnSOD and decrease in complex I abundance at high temperature, or the corresponding decline in condition index, French spat seem to acclimate better to temperature and *p*CO_2_ variation and more precisely adjust their energy metabolism than Norwegian spat. By putatively altering their carbon metabolism to deal with increased redox stress associated with higher temperatures, and by increasing oxygen consumption at elevated *p*CO_2_, potentially to ensure cellular homeostasis, French spat appear better able to maintain growth under OAW conditions. In contrast, Norwegian spat did not appear to fine-tune their proteome, but reduced oxygen consumption if temperature or *p*CO_2_ increased. This corresponded with negative effects on growth, with reduced body weight and shell height when high temperature and *p*CO_2_ were combined.

The experiments were carried out during July, when SST in the Bay of Brest is 2°C higher (16°C) than in the North Sea near the western fjords of Norway (14.0°C; Salt et al. 2016, Omar et al. 2019), which could explain why metabolic rates and growth phenotypes of French spat were not adversely affected by temperature. Furthermore, *p*CO_2_ tends to be higher and more variable in the Bay of Brest (Salt et al. 2016, Omar et al. 2019), and French spat increase their metabolism and maintained growth under elevated *p*CO_2_. However, these metabolic adjustments may be difficult to maintain over longer periods: Harianto et al (2021) found that urchins exposed to elevated *p*CO_2_ and high temperatures after 4 weeks increased metabolism (similar to the French spat in our study), but that after 12 weeks, the combined stress lead to reduced metabolism. The costs of maintaining metabolic function and growth at elevated temperatures could also have contributed towards the reduced survival we observed in French spat. These two populations are known to be genetically divergent (Morvezen et al. 2015), with some genetic differentiation at loci associated with environmental variation in mean annual SST and dissolved organic carbon (Vendrami et al. 2019). This could therefore suggest some adaptive differentiation of these scallop populations in response to environmental variation.

Among marine invertebrate ectotherms, traits as diverse as size (Kelly et al. 2013), metabolic rate (Wood et al. 2016, Osores et al. 2017), developmental plasticity (Pereira et al. 2017), feeding rates (Vargas et al. 2017) and growth (Pespeni et al. 2013a) show important inter-population differences to variation in key environmental variables such as temperature and *p*CO_2_. Our integrative approach helps to disentangle some of the molecular and metabolic differences between populations of this economically important species, highlighting which physiological processes may be involved in acclimatisation processes. Future studies that combine these approaches with genetic studies that estimate the population-specific heritability and plasticity of acclimatory or adaptive traits will be essential in improving our understanding of how populations will respond to global climate change.

## Acknowledgments

We are grateful to Florian Breton from L’Écloserie du Tinduff and to Thorolf Magnesen from Scalpro AS for supplying spat. We thank Julien Thébault for use of the Zeiss Stereomicroscope and Axiovision software. Thanks also to Steven Parratt for discussion about statistical approaches.

## Data availability

The data that support the findings of this study are available to the public at https://github.com/ewan-harney/scallop_oaw

