## Supplementary material for "Impacts of ocean acidification and warming on post-larval growth and metabolism in two populations of the great scallop (*Pecten maximus* L.)"

Figure S1

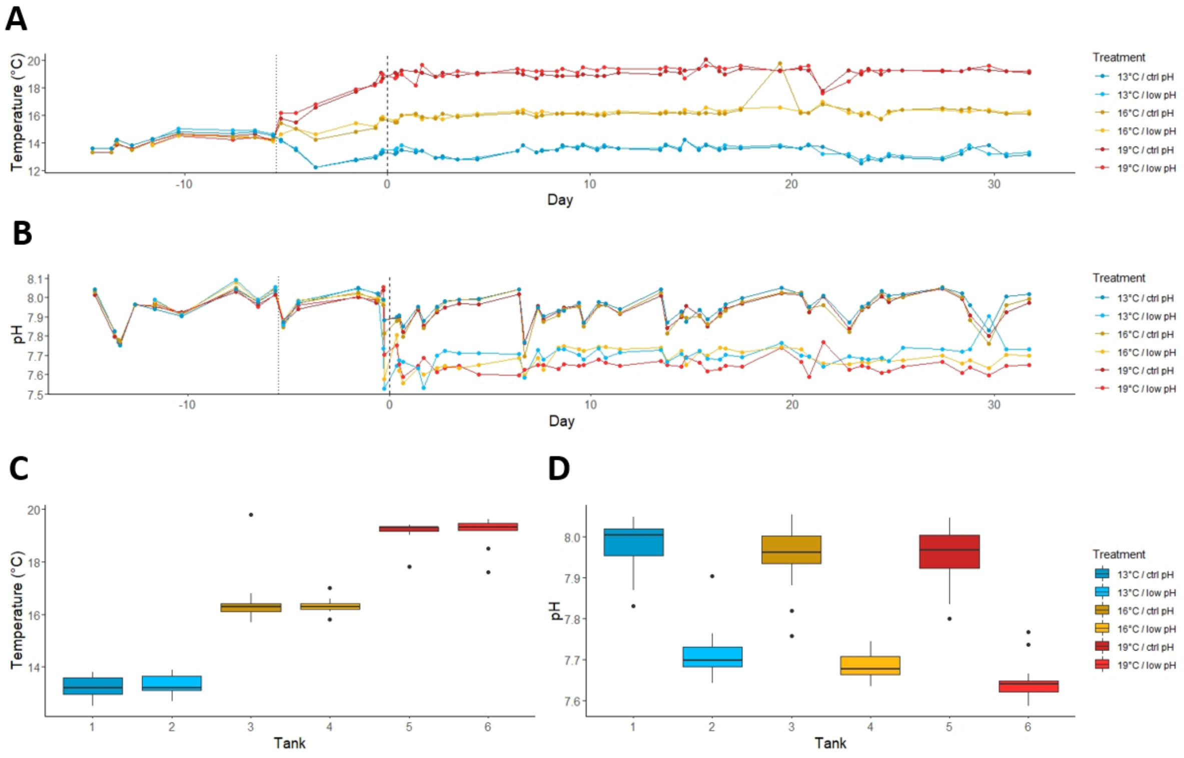


**Fig. S1. Experimental system environmental conditions. A**) Temperatures and **B**) pH for initial acclimation (to dotted line), adjustment period (between dotted and dashed lines) and experiment (from dashed line at day 0 to day 31). Boxplots showing median, interquartile range and whiskers (up to 1.5 × interquartile range) for **C**) temperature and **D**) pH during experimental treatment period (from day 0 to day 31).

Figure S2

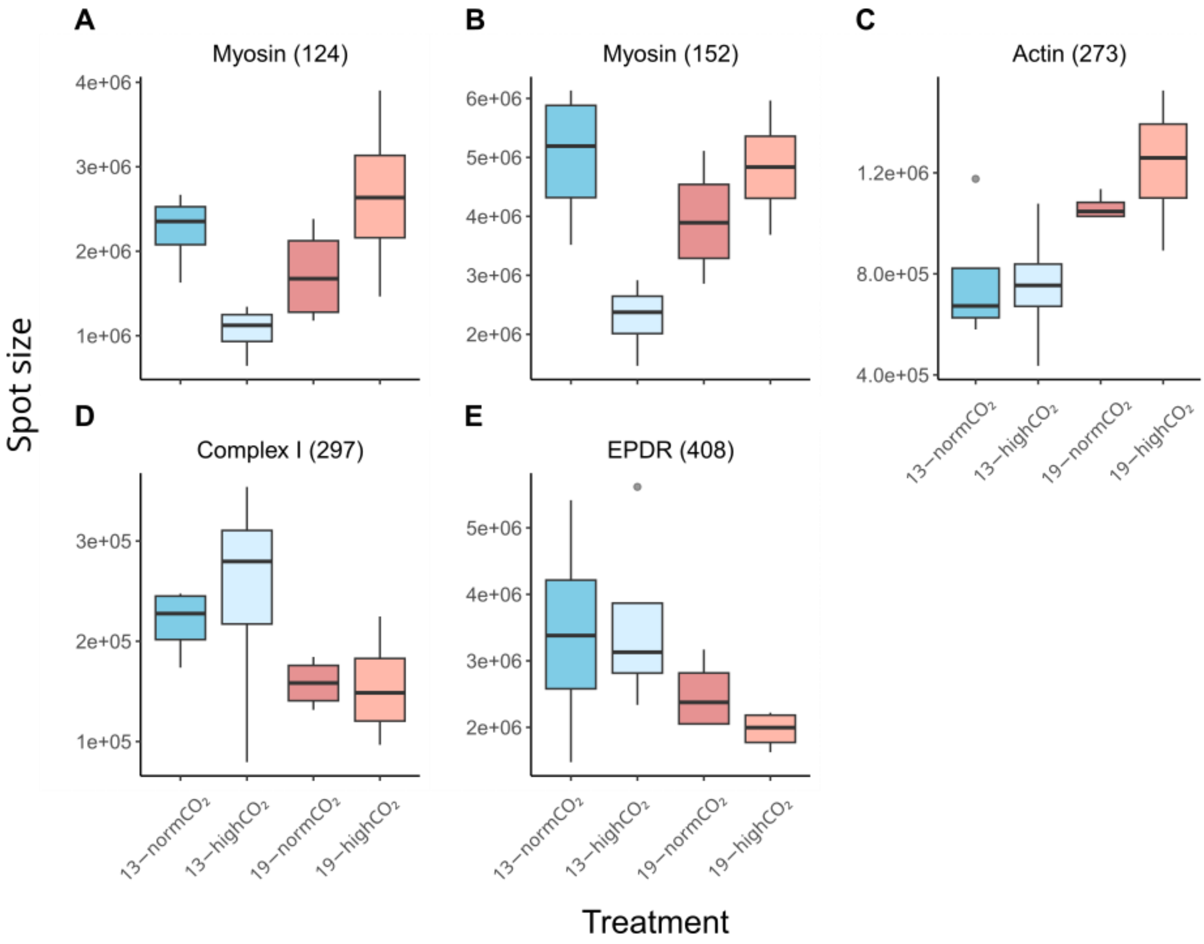


**Fig. S2. Temperature and pH effects on protein abundance in Norwegian spat.** None of the differences were significant at our significance threshold (FDR < 0.05), but the interaction between temperature and pH was significant prior to corrections for multiple testing ($P < 0.01$) in two isoforms of myosin (**A-B**), and at this threshold the temperature effect was significant in an isoform of actin (**C**). Temperature effects on Complex I (**D**) and EPDR (**E**) were also significant at a less stringent uncorrected threshold ($P < 0.05$); comparison with protein abundance patterns in French spat (Fig. 5H and 5K) highlights the weaker patterns of environmental sensitivity in Norwegian spat. For all treatments in all proteins, $n = 4$.

**Table S1.** Contrasts between factor levels for signficant factors in the phenotpyic analysis. Significant contrasts are highlighted in bold

| Trait | Population | Constant | | Contrast | | Estimate | S.E. | DF | t-ratio | P-value |
|---|---|---|---|---|---|---|---|---|---|---|
| Final shell height | Norway | Temp: | 13 | pH: | ctrl - low | -0.281 | 0.240 | 12.9 | -1.172 | 0.2623 |
| | | | 16 | pH: | ctrl - low | 0.249 | 0.236 | 12.4 | 1.055 | 0.3116 |
| | | | **19** | **pH:** | **ctrl - low** | **0.626** | **0.233** | **11.9** | **2.683** | **0.0201** |
| | | pH: | ctrl | Temp: | 13 - 16 | -0.352 | 0.240 | 13.2 | -1.467 | 0.3372 |
| | | | | Temp: | 13 - 19 | -0.307 | 0.236 | 12.4 | -1.299 | 0.4217 |
| | | | | Temp: | 16 - 19 | 0.045 | 0.236 | 12.6 | 0.192 | 0.9799 |
| | | | low | Temp: | 13 - 16 | 0.177 | 0.236 | 12.2 | 0.753 | 0.7375 |
| | | | | Temp: | 13 - 19 | 0.600 | 0.236 | 12.3 | 2.539 | 0.0618 |
| | | | | Temp: | 16 - 19 | 0.422 | 0.232 | 11.7 | 1.817 | 0.2066 |
| Dry body weight | Norway | Temp: | 13 | pH: | ctrl - low | -0.155 | 0.098 | 12.7 | -1.592 | 0.1360 |
| | | | 16 | pH: | ctrl - low | 0.094 | 0.096 | 12.2 | 0.979 | 0.3468 |
| | | | **19** | **pH:** | **ctrl - low** | **0.229** | **0.095** | **11.6** | **2.424** | **0.0328** |
| | | pH: | ctrl | Temp: | 13 - 16 | -0.098 | 0.098 | 13.1 | -1.002 | 0.5885 |
| | | | | Temp: | 13 - 19 | -0.025 | 0.096 | 12.2 | -0.255 | 0.9648 |
| | | | | Temp: | 16 - 19 | 0.073 | 0.096 | 12.4 | 0.764 | 0.7309 |
| | | | low | Temp: | 13 - 16 | 0.151 | 0.096 | 11.9 | 1.579 | 0.2921 |
| | | | | **Temp:** | **13 - 19** | **0.360** | **0.096** | **12.0** | **3.750** | **0.0072** |
| | | | | Temp: | 16 - 19 | 0.209 | 0.094 | 11.4 | 2.218 | 0.1105 |
| Total shell weight | Norway | Temp: | 13 | pH: | ctrl - low | **-0.136** | **0.061** | **12.8** | **-2.241** | **0.0434** |
| | | | 16 | pH: | ctrl - low | 0.093 | 0.059 | 12.3 | 1.562 | 0.1437 |
| | | | 19 | pH: | ctrl - low | 0.124 | 0.059 | 11.7 | 2.120 | 0.0562 |
| | | pH: | ctrl | **Temp:** | **13 - 16** | **-0.162** | **0.061** | **13.2** | **-2.664** | **0.0474** |
| | | | | Temp: | 13 - 19 | -0.139 | 0.060 | 12.3 | -2.339 | 0.0875 |
| | | | | Temp: | 16 - 19 | 0.022 | 0.060 | 12.5 | 0.377 | 0.9253 |
| | | | low | Temp: | 13 - 16 | 0.067 | 0.059 | 12.0 | 1.125 | 0.5178 |
| | | | | Temp: | 13 - 19 | 0.121 | 0.060 | 12.1 | 2.027 | 0.1476 |
| | | | | Temp: | 16 - 19 | 0.054 | 0.058 | 11.5 | 0.924 | 0.6368 |
| Condition index | France | - | - | Temp: | 13 - 16 | 0.003 | 0.002 | 14.5 | 1.850 | 0.1889 |
| | | | | **Temp:** | **13 - 19** | **0.005** | **0.002** | **13.8** | **3.429** | **0.0108** |
| | | | | Temp: | 16 - 19 | 0.002 | 0.002 | 14.6 | 1.535 | 0.3042 |
| | Norway | - | - | Temp: | 13 - 16 | 0.003 | 0.001 | 15.2 | 2.324 | 0.0824 |
| | | | | **Temp:** | **13 - 19** | **0.007** | **0.001** | **14.8** | **5.383** | **0.0002** |
| | | | | **Temp:** | **16 - 19** | **0.004** | **0.001** | **14.6** | **3.069** | **0.0206** |

**Table S2. List of successfully annotated proteins that differ significantly (FDR < 0.05) in response to main effects population, temperature and/or pH.** Log fold change values are highlighted in red (LFC > 1 or FC > 2) or green (LFC > 0.58 or FC > 1.5) and the factor level in which the protein is elevated is provided to aid interpretation. Two proteins (007 and 217) appear twice, due to the presence of two significant main effects.

*(This table is included as a separate excel file)*

**Table S3.** Results of fully factorial linear models for 12 proteins in which two or more main effects or interactions were found to be significant (FDR < 0.05). Significant FDR values are highlighted in red and the factor level (or combination of levels) in which the protein is elevated is provided to aid interpretation.

| spot | Accession | Description | Species | Effect/Interaction | Elevated in... | df | SumSq | MeanSq | Fval | Pval | fdr |
|---|---|---|---|---|---|---|---|---|---|---|---|
| sp_196 | OWF48408.1 | Retinal dehydrogenase 2 | *Mizuhopecten yessoensis* | Population | French | 1 | 3.11E+11 | 3.11E+11 | 28.7206 | 1.68E-05 | 0.0003 |
| sp_196 | OWF48408.1 | Retinal dehydrogenase 2 | *Mizuhopecten yessoensis* | Temp | | 1 | 6.64E+10 | 6.64E+10 | 6.1293 | 0.020742 | 0.1092 |
| sp_196 | OWF48408.1 | Retinal dehydrogenase 2 | *Mizuhopecten yessoensis* | pH | | 1 | 1.10E+10 | 1.10E+10 | 1.0173 | 0.323227 | 0.6250 |
| sp_196 | OWF48408.1 | Retinal dehydrogenase 2 | *Mizuhopecten yessoensis* | Population:Temp | French:19C | 1 | 1.66E+11 | 1.66E+11 | 15.3289 | 0.000653 | 0.0060 |
| sp_196 | OWF48408.1 | Retinal dehydrogenase 2 | *Mizuhopecten yessoensis* | Population:pH | | 1 | 4.20E+10 | 4.20E+10 | 3.8804 | 0.060498 | 0.2497 |
| sp_196 | OWF48408.1 | Retinal dehydrogenase 2 | *Mizuhopecten yessoensis* | Temp:pH | | 1 | 2.38E+10 | 2.38E+10 | 2.1993 | 0.151086 | 0.4306 |
| sp_196 | OWF48408.1 | Retinal dehydrogenase 2 | *Mizuhopecten yessoensis* | Population:Temp:pH | | 1 | 1.86E+10 | 1.86E+10 | 1.7144 | 0.202808 | 0.4941 |
| sp_273 | OWF37106.1 | Actin, cytoplasmic | *Mizuhopecten yessoensis* | Population | Norwegian | 1 | 3.77E+12 | 3.77E+12 | 131.0622 | 3.29E-11 | 0.0000 |
| sp_273 | OWF37106.1 | Actin, cytoplasmic | *Mizuhopecten yessoensis* | Temp | 19C | 1 | 3.09E+11 | 3.09E+11 | 10.7291 | 0.003197 | 0.0236 |
| sp_273 | OWF37106.1 | Actin, cytoplasmic | *Mizuhopecten yessoensis* | pH | | 1 | 1.69E+10 | 1.69E+10 | 0.5864 | 0.451267 | 0.7304 |
| sp_273 | OWF37106.1 | Actin, cytoplasmic | *Mizuhopecten yessoensis* | Population:Temp | Norwegian:19C | 1 | 2.79E+11 | 2.79E+11 | 9.6873 | 0.004744 | 0.0316 |
| sp_273 | OWF37106.1 | Actin, cytoplasmic | *Mizuhopecten yessoensis* | Population:pH | | 1 | 6.94E+09 | 6.94E+09 | 0.241 | 0.62795 | 0.8348 |
| sp_273 | OWF37106.1 | Actin, cytoplasmic | *Mizuhopecten yessoensis* | Temp:pH | | 1 | 3.40E+10 | 3.40E+10 | 1.1804 | 0.28806 | 0.5966 |
| sp_273 | OWF37106.1 | Actin, cytoplasmic | *Mizuhopecten yessoensis* | Population:Temp:pH | | 1 | 7.18E+09 | 7.18E+09 | 0.2494 | 0.622076 | 0.8347 |
| sp_239 | Q26065.1 | Actin, adductor muscle | *Placopecten magellanicus* | Population | French | 1 | 1.43E+13 | 1.43E+13 | 12.3384 | 0.001786 | 0.0145 |
| sp_239 | Q26065.1 | Actin, adductor muscle | *Placopecten magellanicus* | Temp | | 1 | 1.51E+12 | 1.51E+12 | 1.3058 | 0.264429 | 0.5665 |
| sp_239 | Q26065.1 | Actin, adductor muscle | *Placopecten magellanicus* | pH | | 1 | 8.32E+12 | 8.32E+12 | 7.1848 | 0.013079 | 0.0738 |
| sp_239 | Q26065.1 | Actin, adductor muscle | *Placopecten magellanicus* | Population:Temp | | 1 | 7.99E+11 | 7.99E+11 | 0.6899 | 0.414375 | 0.6965 |
| sp_239 | Q26065.1 | Actin, adductor muscle | *Placopecten magellanicus* | Population:pH | French:pH7.7 | 1 | 1.05E+13 | 1.05E+13 | 9.1043 | 0.005955 | 0.0387 |
| sp_239 | Q26065.1 | Actin, adductor muscle | *Placopecten magellanicus* | Temp:pH | | 1 | 2.64E+12 | 2.64E+12 | 2.2791 | 0.144181 | 0.4245 |
| sp_239 | Q26065.1 | Actin, adductor muscle | *Placopecten magellanicus* | Population:Temp:pH | | 1 | 1.22E+12 | 1.22E+12 | 1.058 | 0.313917 | 0.6116 |
| sp_419 | XP_021346372.1 | triosephosphate isomerase B-like | *Mizuhopecten yessoensis* | Population | French | 1 | 5.89E+11 | 5.89E+11 | 24.9958 | 4.16E-05 | 0.0006 |
| sp_419 | XP_021346372.1 | triosephosphate isomerase B-like | *Mizuhopecten yessoensis* | Temp | | 1 | 1.83E+11 | 1.83E+11 | 7.7552 | 0.01029 | 0.0599 |
| sp_419 | XP_021346372.1 | triosephosphate isomerase B-like | *Mizuhopecten yessoensis* | pH | | 1 | 7.14E+10 | 7.14E+10 | 3.0332 | 0.09438 | 0.3389 |
| sp_419 | XP_021346372.1 | triosephosphate isomerase B-like | *Mizuhopecten yessoensis* | Population:Temp | | 1 | 1.37E+11 | 1.37E+11 | 5.7991 | 0.02408 | 0.1233 |
| sp_419 | XP_021346372.1 | triosephosphate isomerase B-like | *Mizuhopecten yessoensis* | Population:pH | French:pH8.0 | 1 | 3.01E+11 | 3.01E+11 | 12.7984 | 0.00152 | 0.0127 |
| sp_419 | XP_021346372.1 | triosephosphate isomerase B-like | *Mizuhopecten yessoensis* | Temp:pH | | 1 | 4.95E+10 | 4.95E+10 | 2.101 | 0.16015 | 0.4428 |
| sp_419 | XP_021346372.1 | triosephosphate isomerase B-like | *Mizuhopecten yessoensis* | Population:Temp:pH | | 1 | 5.56E+10 | 5.56E+10 | 2.363 | 0.13733 | 0.4150 |
| sp_124 | AAD52842.1 | myosin heavy chain | *Pecten maximus* | Population | Norwegian | 1 | 6.57E+12 | 6.57E+12 | 24.9507 | 4.21E-05 | 0.0006 |
| sp_124 | AAD52842.1 | myosin heavy chain | *Pecten maximus* | Temp | | 1 | 6.69E+11 | 6.69E+11 | 2.5405 | 0.124042 | 0.3921 |

| | | | | | | | | | | | |
|---|---|---|---|---|---|---|---|---|---|---|---|
| sp_124 | AAD52842.1 | myosin heavy chain | *Pecten maximus* | pH | | 1 | 2.50E+08 | 2.50E+08 | 0.0009 | 0.97569 | 0.9955 |
| sp_124 | AAD52842.1 | myosin heavy chain | *Pecten maximus* | Population:Temp | | 1 | 4.98E+11 | 4.98E+11 | 1.8884 | 0.182083 | 0.4619 |
| sp_124 | AAD52842.1 | myosin heavy chain | *Pecten maximus* | Population:pH | | 1 | 1.48E+11 | 1.48E+11 | 0.5612 | 0.461058 | 0.7349 |
| sp_124 | AAD52842.1 | myosin heavy chain | *Pecten maximus* | Temp:pH | 13C:pH8.0 / 19C:pH7.7 | 1 | 4.72E+12 | 4.72E+12 | 17.909 | 0.000293 | 0.0031 |
| sp_124 | AAD52842.1 | myosin heavy chain | *Pecten maximus* | Population:Temp:pH | | 1 | 6.92E+11 | 6.92E+11 | 2.6265 | 0.118159 | 0.3844 |
| sp_290 | Q26065.1 | Actin, adductor muscle | *Placopecten magellanicus* | Population | Norwegian | 1 | 8.49E+13 | 8.49E+13 | 21.9959 | 9.13E-05 | 0.0012 |
| sp_290 | Q26065.1 | Actin, adductor muscle | *Placopecten magellanicus* | Temp | | 1 | 4.11E+12 | 4.11E+12 | 1.0648 | 0.312406 | 0.6116 |
| sp_290 | Q26065.1 | Actin, adductor muscle | *Placopecten magellanicus* | pH | | 1 | 1.67E+13 | 1.67E+13 | 4.3343 | 0.048185 | 0.2202 |
| sp_290 | Q26065.1 | Actin, adductor muscle | *Placopecten magellanicus* | Population:Temp | | 1 | 3.58E+12 | 3.58E+12 | 0.9274 | 0.345131 | 0.6362 |
| sp_290 | Q26065.1 | Actin, adductor muscle | *Placopecten magellanicus* | Population:pH | | 1 | 2.74E+12 | 2.74E+12 | 0.7096 | 0.407886 | 0.6919 |
| sp_290 | Q26065.1 | Actin, adductor muscle | *Placopecten magellanicus* | Temp:pH | 13C:pH8.0 / 19C:pH7.7 | 1 | 3.36E+13 | 3.36E+13 | 8.7196 | 0.006939 | 0.0417 |
| sp_290 | Q26065.1 | Actin, adductor muscle | *Placopecten magellanicus* | Population:Temp:pH | | 1 | 1.14E+12 | 1.14E+12 | 0.2968 | 0.59095 | 0.8129 |
| sp_152 | AAD52842.1 | myosin heavy chain | *Pecten maximus* | Population | Norwegian | 1 | 2.51E+13 | 2.51E+13 | 36.4406 | 3.11E-06 | 0.0001 |
| sp_152 | AAD52842.1 | myosin heavy chain | *Pecten maximus* | Temp | 19C | 1 | 6.32E+12 | 6.32E+12 | 9.1662 | 0.005812 | 0.0383 |
| sp_152 | AAD52842.1 | myosin heavy chain | *Pecten maximus* | pH | | 1 | 1.53E+11 | 1.53E+11 | 0.2223 | 0.641573 | 0.8414 |
| sp_152 | AAD52842.1 | myosin heavy chain | *Pecten maximus* | Population:Temp | | 1 | 1.80E+11 | 1.80E+11 | 0.2613 | 0.61387 | 0.8300 |
| sp_152 | AAD52842.1 | myosin heavy chain | *Pecten maximus* | Population:pH | | 1 | 4.85E+12 | 4.85E+12 | 7.0323 | 0.013961 | 0.0780 |
| sp_152 | AAD52842.1 | myosin heavy chain | *Pecten maximus* | Temp:pH | 13C:pH8.0 / 19C:pH7.7 | 1 | 1.63E+13 | 1.63E+13 | 23.6562 | 5.87E-05 | 0.0008 |
| sp_152 | AAD52842.1 | myosin heavy chain | *Pecten maximus* | Population:Temp:pH | | 1 | 1.16E+12 | 1.16E+12 | 1.6903 | 0.205906 | 0.4951 |
| sp_442 | XP_021365581.1 | glutathione S-transferase P-like | *Mizuhopecten yessoensis* | Population | French | 1 | 1.70E+12 | 1.70E+12 | 26.7172 | 2.71E-05 | 0.0005 |
| sp_442 | XP_021365581.1 | glutathione S-transferase P-like | *Mizuhopecten yessoensis* | Temp | | 1 | 2.57E+11 | 2.57E+11 | 4.0463 | 0.055626 | 0.2341 |
| sp_442 | XP_021365581.1 | glutathione S-transferase P-like | *Mizuhopecten yessoensis* | pH | | 1 | 1.48E+11 | 1.48E+11 | 2.3374 | 0.139378 | 0.4166 |
| sp_442 | XP_021365581.1 | glutathione S-transferase P-like | *Mizuhopecten yessoensis* | Population:Temp | | 1 | 8.83E+10 | 8.83E+10 | 1.3909 | 0.249819 | 0.5593 |
| sp_442 | XP_021365581.1 | glutathione S-transferase P-like | *Mizuhopecten yessoensis* | Population:pH | | 1 | 7.31E+10 | 7.31E+10 | 1.1519 | 0.293826 | 0.6018 |
| sp_442 | XP_021365581.1 | glutathione S-transferase P-like | *Mizuhopecten yessoensis* | Temp:pH | | 1 | 1.57E+11 | 1.57E+11 | 2.4762 | 0.128671 | 0.3979 |
| sp_442 | XP_021365581.1 | glutathione S-transferase P-like | *Mizuhopecten yessoensis* | Population:Temp:pH | French:19C:pH8.0 | 1 | 5.26E+11 | 5.26E+11 | 8.2871 | 0.008263 | 0.0491 |
| sp_447 | XP_021374826.1 | unch prot LOC110464099 | *Mizuhopecten yessoensis* | Population | French | 1 | 3.20E+12 | 3.20E+12 | 97.645 | 6.20E-10 | 0.0000 |
| sp_447 | XP_021374826.1 | unch prot LOC110464100 | *Mizuhopecten yessoensis* | Temp | 19C | 1 | 3.31E+11 | 3.31E+11 | 10.0971 | 0.004055 | 0.0277 |
| sp_447 | XP_021374826.1 | unch prot LOC110464101 | *Mizuhopecten yessoensis* | pH | | 1 | 1.28E+11 | 1.28E+11 | 3.9161 | 0.059412 | 0.2470 |
| sp_447 | XP_021374826.1 | unch prot LOC110464102 | *Mizuhopecten yessoensis* | Population:Temp | | 1 | 9.98E+10 | 9.98E+10 | 3.0438 | 0.09384 | 0.3389 |
| sp_447 | XP_021374826.1 | unch prot LOC110464103 | *Mizuhopecten yessoensis* | Population:pH | Fr:pH8.0 / No:pH7.7 | 1 | 5.97E+11 | 5.97E+11 | 18.2055 | 0.000268 | 0.0029 |
| sp_447 | XP_021374826.1 | unch prot LOC110464104 | *Mizuhopecten yessoensis* | Temp:pH | | 1 | 4.36E+10 | 4.36E+10 | 1.3314 | 0.259916 | 0.5659 |
| sp_447 | XP_021374826.1 | unch prot LOC110464105 | *Mizuhopecten yessoensis* | Population:Temp:pH | | 1 | 1.47E+10 | 1.47E+10 | 0.4498 | 0.508851 | 0.7688 |

**Table S4**: Results of post hoc tests comparing mean protein abundance for all significantly differentially abundant proteins among French spat. Significant contrasts are highlighted in bold.

| Protein | Description | Contrast | Estimate | S.E. | DF | t ratio | adj. P val |
|---|---|---|---|---|---|---|---|
| sp_144 | myosin heavy chain | 13°C pH 8.0 - 13°C pH 7.7 | -248696 | 94953 | 12 | -2.619 | 0.0907 |
| | | 13°C pH 8.0 - 19°C pH 8.0 | 126971 | 94953 | 12 | 1.337 | 0.5585 |
| | | 13°C pH 8.0 - 19°C pH 7.7 | -276952 | 94953 | 12 | -2.917 | 0.0547 |
| | | **13°C pH 7.7 - 19°C pH 8.0** | **375667** | **94953** | **12** | **3.956** | **0.0089** |
| | | 13°C pH 7.7 - 19°C pH 7.7 | -28256 | 94953 | 12 | -0.298 | 0.9904 |
| | | **19°C pH 8.0 - 19°C pH 7.7** | **-403923** | **94953** | **12** | **-4.254** | **0.0053** |
| sp_196 | Retinal dehydrogenase 2 | 13°C pH 8.0 - 13°C pH 7.7 | 6849 | 72050 | 12 | 0.095 | 0.9997 |
| | | **13°C pH 8.0 - 19°C pH 8.0** | **-337966** | **72050** | **12** | **-4.691** | **0.0025** |
| | | 13°C pH 8.0 - 19°C pH 7.7 | -125586 | 72050 | 12 | -1.743 | 0.3452 |
| | | **13°C pH 7.7 - 19°C pH 8.0** | **-344815** | **72050** | **12** | **-4.786** | **0.0022** |
| | | 13°C pH 7.7 - 19°C pH 7.7 | -132435 | 72050 | 12 | -1.838 | 0.3036 |
| | | 19°C pH 8.0 - 19°C pH 7.7 | 212379 | 72050 | 12 | 2.948 | 0.0519 |
| sp_217 | ATP synthase subunit beta, mitochondrial | 13°C pH 8.0 - 13°C pH 7.7 | 4048976 | 1529768 | 12 | 2.647 | 0.0866 |
| | | 13°C pH 8.0 - 19°C pH 8.0 | -3642971 | 1529768 | 12 | -2.381 | 0.134 |
| | | 13°C pH 8.0 - 19°C pH 7.7 | 810992 | 1529768 | 12 | 0.53 | 0.9501 |
| | | **13°C pH 7.7 - 19°C pH 8.0** | **-7691946** | **1529768** | **12** | **-5.028** | **0.0015** |
| | | 13°C pH 7.7 - 19°C pH 7.7 | -3237984 | 1529768 | 12 | -2.117 | 0.2027 |
| | | 19°C pH 8.0 - 19°C pH 7.7 | 4453963 | 1529768 | 12 | 2.912 | 0.0552 |
| sp_252 | medium-chain specific acyl-CoA dehydrogenase, mitochondrial-like | 13°C pH 8.0 - 13°C pH 7.7 | -331168 | 113480 | 12 | -2.918 | 0.0546 |
| | | **13°C pH 8.0 - 19°C pH 8.0** | **347428** | **113480** | **12** | **3.062** | **0.0426** |
| | | 13°C pH 8.0 - 19°C pH 7.7 | 78985 | 113480 | 12 | 0.696 | 0.8966 |
| | | **13°C pH 7.7 - 19°C pH 8.0** | **678596** | **113480** | **12** | **5.98** | **0.0003** |
| | | **13°C pH 7.7 - 19°C pH 7.7** | **410153** | **113480** | **12** | **3.614** | **0.0162** |
| | | 19°C pH 8.0 - 19°C pH 7.7 | -268443 | 113480 | 12 | -2.366 | 0.1374 |
| sp_297 | complex I | 13°C pH 8.0 - 13°C pH 7.7 | 95460 | 32233 | 12 | 2.962 | 0.0506 |
| | | **13°C pH 8.0 - 19°C pH 8.0** | **120616** | **32233** | **12** | **3.742** | **0.013** |
| | | **13°C pH 8.0 - 19°C pH 7.7** | **157709** | **32233** | **12** | **4.893** | **0.0018** |
| | | 13°C pH 7.7 - 19°C pH 8.0 | 25157 | 32233 | 12 | 0.78 | 0.8618 |
| | | 13°C pH 7.7 - 19°C pH 7.7 | 62249 | 32233 | 12 | 1.931 | 0.2664 |
| | | 19°C pH 8.0 - 19°C pH 7.7 | 37093 | 32233 | 12 | 1.151 | 0.667 |
| sp_356 | Actin, cytoplasmic | 13°C pH 8.0 - 13°C pH 7.7 | 223717 | 178836 | 12 | 1.251 | 0.6086 |
| | | 13°C pH 8.0 - 19°C pH 8.0 | -491499 | 178836 | 12 | -2.748 | 0.0729 |
| | | 13°C pH 8.0 - 19°C pH 7.7 | -272767 | 178836 | 12 | -1.525 | 0.4536 |
| | | **13°C pH 7.7 - 19°C pH 8.0** | **-715216** | **178836** | **12** | **-3.999** | **0.0083** |
| | | 13°C pH 7.7 - 19°C pH 7.7 | -496484 | 178836 | 12 | -2.776 | 0.0696 |
| | | 19°C pH 8.0 - 19°C pH 7.7 | 218732 | 178836 | 12 | 1.223 | 0.6249 |
| sp_408 | mammalian ependymin-related protein 1-like | 13°C pH 8.0 - 13°C pH 7.7 | -832114 | 496224 | 12 | -1.677 | 0.3763 |
| | | 13°C pH 8.0 - 19°C pH 8.0 | 1373654 | 496224 | 12 | 2.768 | 0.0705 |
| | | **13°C pH 8.0 - 19°C pH 7.7** | **1388933** | **496224** | **12** | **2.799** | **0.0669** |
| | | **13°C pH 7.7 - 19°C pH 8.0** | **2205767** | **496224** | **12** | **4.445** | **0.0038** |
| | | **13°C pH 7.7 - 19°C pH 7.7** | **2221047** | **496224** | **12** | **4.476** | **0.0036** |
| | | 19°C pH 8.0 - 19°C pH 7.7 | 15280 | 496224 | 12 | 0.031 | 1 |
| sp_415 | triosephosphate isomerase B-like | 13°C pH 8.0 - 13°C pH 7.7 | 336339 | 138339 | 12 | 2.431 | 0.1236 |
| | | 13°C pH 8.0 - 19°C pH 8.0 | -311134 | 138339 | 12 | -2.249 | 0.1653 |
| | | 13°C pH 8.0 - 19°C pH 7.7 | -127134 | 138339 | 12 | -0.919 | 0.7954 |
| | | **13°C pH 7.7 - 19°C pH 8.0** | **-647473** | **138339** | **12** | **-4.68** | **0.0026** |

| | | | | | | | |
|---|---|---|---|---|---|---|---|
| | | **13°C pH 7.7 - 19°C pH 7.7** | **-463473** | **138339** | **12** | **-3.35** | **0.0257** |
| | | 19°C pH 8.0 - 19°C pH 7.7 | 184001 | 138339 | 12 | 1.33 | 0.5626 |
| sp_447 | uncharacterized protein LOC110464099 (KY) | 13°C pH 8.0 - 13°C pH 7.7 | 283002 | 145810 | 12 | 1.941 | 0.2627 |
| | | 13°C pH 8.0 - 19°C pH 8.0 | -431863 | 145810 | 12 | -2.962 | 0.0506 |
| | | 13°C pH 8.0 - 19°C pH 7.7 | 84714 | 145810 | 12 | 0.581 | 0.9359 |
| | | **13°C pH 7.7 - 19°C pH 8.0** | **-714866** | **145810** | **12** | **-4.903** | **0.0018** |
| | | 13°C pH 7.7 - 19°C pH 7.7 | -198288 | 145810 | 12 | -1.36 | 0.5455 |
| | | **19°C pH 8.0 - 19°C pH 7.7** | **516578** | **145810** | **12** | **3.543** | **0.0184** |
| sp_462 | peptidyl-prolyl cis-trans isomerase B-like | 13°C pH 8.0 - 13°C pH 7.7 | 279276 | 94093 | 12 | 2.968 | 0.0501 |
| | | **13°C pH 8.0 - 19°C pH 8.0** | **495974** | **94093** | **12** | **5.271** | **0.001** |
| | | **13°C pH 8.0 - 19°C pH 7.7** | **677353** | **94093** | **12** | **7.199** | **0.0001** |
| | | 13°C pH 7.7 - 19°C pH 8.0 | 216699 | 94093 | 12 | 2.303 | 0.1518 |
| | | **13°C pH 7.7 - 19°C pH 7.7** | **398077** | **94093** | **12** | **4.231** | **0.0055** |
| | | 19°C pH 8.0 - 19°C pH 7.7 | 181378 | 94093 | 12 | 1.928 | 0.2677 |
| sp_463 | Mn SOD | 13°C pH 8.0 - 13°C pH 7.7 | 21553 | 35064 | 12 | 0.615 | 0.9254 |
| | | **13°C pH 8.0 - 19°C pH 8.0** | **-119923** | **35064** | **12** | **-3.42** | **0.0228** |
| | | **13°C pH 8.0 - 19°C pH 7.7** | **-108786** | **35064** | **12** | **-3.103** | **0.0397** |
| | | **13°C pH 7.7 - 19°C pH 8.0** | **-141477** | **35064** | **12** | **-4.035** | **0.0078** |
| | | **13°C pH 7.7 - 19°C pH 7.7** | **-130340** | **35064** | **12** | **-3.717** | **0.0135** |
| | | 19°C pH 8.0 - 19°C pH 7.7 | 11137 | 35064 | 12 | 0.318 | 0.9883 |
| sp_468 | uncharacterized protein LOC110453073 (vWA) | 13°C pH 8.0 - 13°C pH 7.7 | -165706 | 511581 | 12 | -0.324 | 0.9877 |
| | | **13°C pH 8.0 - 19°C pH 8.0** | **-2442993** | **511581** | **12** | **-4.775** | **0.0022** |
| | | 13°C pH 8.0 - 19°C pH 7.7 | -851666 | 511581 | 12 | -1.665 | 0.3822 |
| | | **13°C pH 7.7 - 19°C pH 8.0** | **-2277287** | **511581** | **12** | **-4.451** | **0.0038** |
| | | 13°C pH 7.7 - 19°C pH 7.7 | -685960 | 511581 | 12 | -1.341 | 0.5564 |
| | | **19°C pH 8.0 - 19°C pH 7.7** | **1591327** | **511581** | **12** | **3.111** | **0.0391** |
